# Hematopoietic recovery after transplantation is primarily derived from the stochastic contribution of hematopoietic stem cells

**DOI:** 10.1101/2021.09.21.461235

**Authors:** Stefan Radtke, Mark Enstrom, Dnyanada Pande, Margaret Cui, Ravishankar Madhu, Anai M. Perez, Hans-Peter Kiem

**Affiliations:** Stem Cell and Gene Therapy Program, Clinical Research Division, Fred Hutchinson Cancer Research Center, Seattle, WA, 98109, USA; Department of Medicine, University of Washington School of Medicine, Seattle, WA, 98195, USA; Department of Pathology, University of Washington School of Medicine, Seattle, WA, 98195, USA

**Keywords:** Hematopoietic stem cells (HSCs), Retroviral integration site analysis, HSC Clone Tracking, Hematopoietic reconstitution, HSC transplantation, Stochastic HSC engraftment

## Abstract

Reconstitution after hematopoietic stem cell (HSC) transplantation is assumed to occur in two distinct phases: initial recovery mediated by short-term progenitors and long-term repopulation by multipotent HSCs which do not contribute to hematopoietic reconstitution during the first 6-9 months. We have previously reported the transplantation and exclusive engraftment of the HSC-enriched CD34^+^CD45RA^-^CD90^+^ phenotype in a nonhuman primate model. Here, we closely followed the clonal diversity and kinetics in these animals. Enhanced sampling and high density clonal tracking within the first 3 month revealed that multipotent HSCs actively contributed to the early phases of neutrophil recovery and became the dominant source for blood cells as early as 50 days after transplant. Longitudinal changes in clonal diversity supported a stochastic engraftment of HSCs with the majority of HSCs clones vanishing early during neutrophil recovery and a smaller fraction of HSC clones expanding into bigger pools to support long-term hematopoiesis. In contrast to the bi-phasic model, we propose that hematopoietic recovery after myeloablation and transplantation is primarily derived from HSCs in a stochastic manner rather than in two phases by independent cell populations.

## INTRODUCTION

Hematopoietic recovery after myeloablative conditioning and hematopoietic stem cell (HSC) transplantation is thought to occur in two distinct waves, with short-term engrafting hematopoietic progenitor cells driving the recovery within the first 6-9 months and long-term engrafting multipotent HSCs providing long-term repopulation^1^. This bi-phasic model has been presumed for several decades, with more recent clonal tracking data from autologous human gene therapy trials and nonhuman primate (NHP) transplantation studies seeming to confirm this concept in human hematopoiesis after transplantation^2-4^. However, the temporal involvement of HSCs after transplant still remains subject to controversy. The use of diverse tracing methods, species-specific differences, as well as variations in the modelling approaches further complicate the comparison and interpretation of data.

Most recent reports simulating the contribution of HSCs in patients are based on the longitudinal tracking of thousands of gene-marked cells using retroviral integration site analysis (ISA)^2,5^. ISA is routinely used to follow gene therapy patients and monitor the potential outgrowth of dominant or malignant clones after gene therapy. While this technology is very reliable in detecting highly abundant clones, low sensitivity and high error rates require significant data exclusion and sophisticated statistical tests to ensure data reliability^2,5,6^. Lack of sensitivity can be overcome by increasing the frequency of sampling (high density) as well as repeated sampling^5,6^. However, limited material from patients remains a bottleneck for improved data quality and, consequently, correct interpretation of such complex datasets.

Our recent studies in the nonhuman primate (NHP) large animal model suggested significantly earlier contribution of multipotent hematopoietic stem and progenitor cells (HSPCs) after autologous transplantation of gene-marked cells than proposed in the bi-phasic model^7^. Here, we performed high-density sampling for ISA in NHPs after transplantation of gene-modified HSCs to overcome the limitations of ISA, reliably determine the onset of HSC contribution during hematopoietic recovery, and closely investigate the kinetics of hematopoietic recovery. Especially during the first month of hematopoietic recovery weekly blood samples were taken in order to enhance ISA data density, increase the reliability to detect clones with low abundance, and determine the kinetics of hematopoietic reconstitution during neutrophil recovery. Animals were followed for 4.5 years to confirm that tracked HSC clones persist long-term. Integration site (IS) profiles were compared to historic control animals transplanted with bulk CD34^+^ cells to rule out an impact of the experimental design on the clonal tracking. Finally, observed changes in clonal diversity *in vivo* were used to inform a simulation of hematopoietic reconstitution, determine the temporal involvement of HSCs, and refine the phases of hematopoietic recovery after myeloablation and HSC transplantation.

## RESULTS

### Bone marrow recovery and long-term multilineage engraftment are primarily driven by CD34^+^CD90^+^ cells

Peripheral blood (PB) from animals transplanted with gene-modified CD34 subsets (**Figure 1A**) ^7^ was analyzed flow-cytometrically and white blood cells (WBCs) were collected for ISA (**Figure 1B**). Fluorochrome-expressing lineages were FACS (fluorescence assisted cell sorting)-purified for ISA from Z13264 and Z14004 (**Figure 1B**, black symbols). Due to low gene marking and limited blood volume, bulk lineages containing 2-5% fluorochrome-expressing cells were isolated for Z15086. Bone marrow (BM) was analyzed flow-cytometrically and CD34^+^ cells isolated for functional assays as well as ISA (**Figure 1B**).

**Figure 1.**
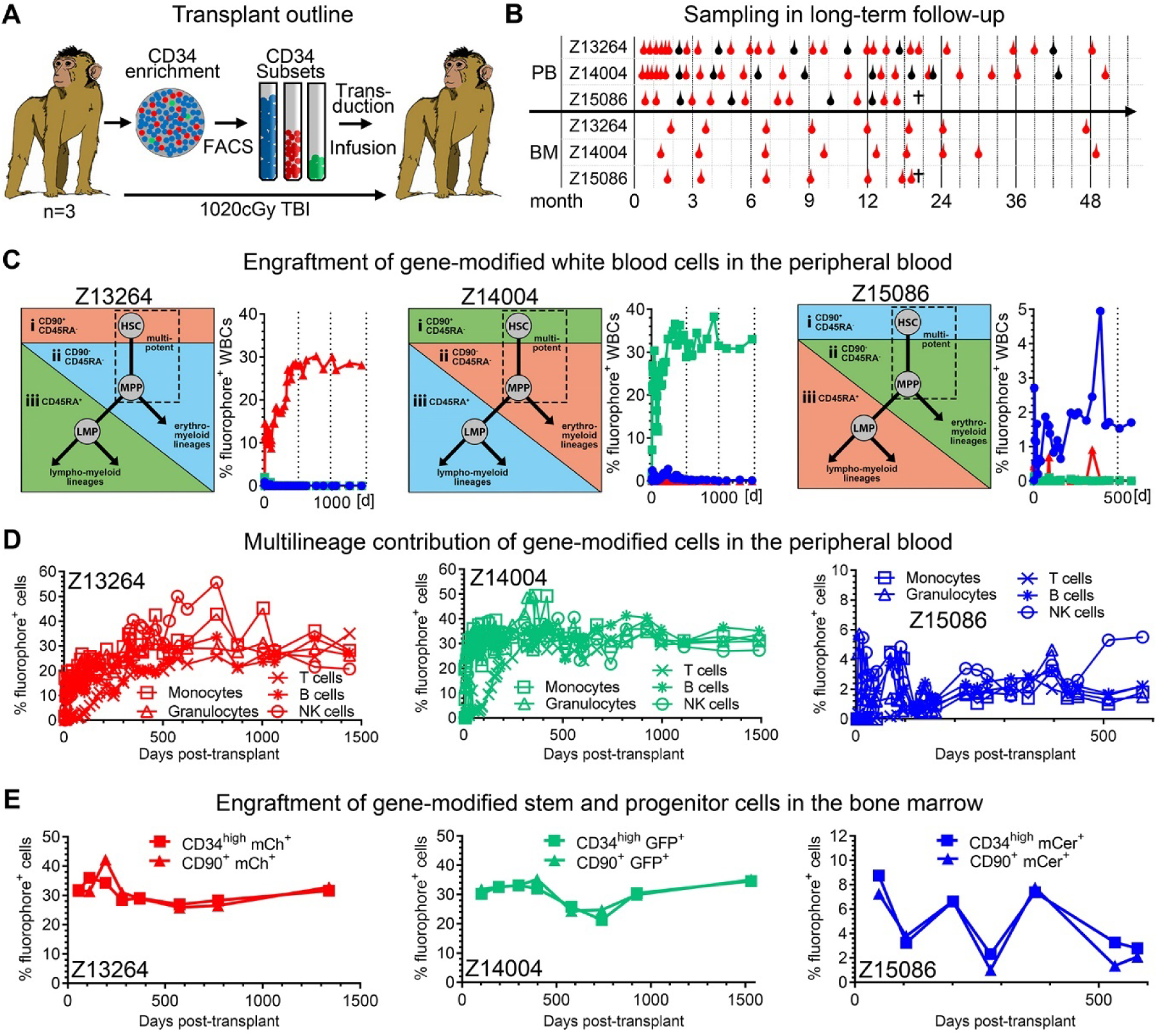
Longitudinal assessment of PB and BM after competitive transplantation of gene-modified CD34 subsets. (**A**) CD34^+^ HSPCs from three pigtail macaques were enriched with immunomagnetic beads, CD34 subsets (CD90^+^CD45RA^−^, CD90^-^CD45RA^−^, and CD90^−^CD45RA^+^) FACS-purified, transduced, and co-transplanted after myeloablative conditioning with total body irradiation (TBI). **(B)** Animals were followed for up to 50 months taking PB and BM draws. Bulk WBCs were collected as indicated with red symbols, whereas blood lineages (T cells: CD3^+^, B cells: CD20^+^, NK cells: CD16^+^, Monocytes: CD14^+^, Granulocytes: CD11b^+^CD14^-^) were FACS-purified as indicated with black symbols. **(C-E)** Expression of fluorochromes in PB WBCs (C), PB lineages (D), and BM CD34^+^/CD34^+^CD90^+^ HSPCs (E) was longitudinally tracked by flow cytometry.

Longitudinal flow-cytometric analysis confirmed that CD34^+^CD45RA^-^CD90^+^ cells remained the exclusive source of all mature blood cells in the PB throughout the entire follow-up (**Figure 1C,D**). Gene marking in PB WBCs continuously increased within the first 9 to 12 months, reaching a plateau thereafter. Simultaneously, WBC, lymphocyte, red blood cell (RBC), and platelet counts stabilized within normal range and remained stable (**Figure S1A**). Nearly identical gene marking efficiency was found in T cells, B cells, NK cells, granulocytes and monocytes starting at 9 months post-transplant (**Figure 1D**).

Gene marking of CD34^+^ hematopoietic stem and progenitor cells (HSPCs), as well as the HSC-enriched CD34^+^CD45RA^-^CD90^+^ subset in the BM, mirrored observed frequencies in the PB (**Figure 1E**). High gene marking was observed significantly earlier in the BM stem cell compartment compared to PB WBCs, reaching stable levels of gene-modification by week 6 (**Figure S1B**). Gene-modified CD34^+^ and CD34^+^CD45RA^−^CD90^+^ demonstrated almost identical erythro-myeloid colony-forming potential in comparison to their unmarked counterparts, indicating no impact of FACS-purification and *ex vivo* gene-modification on their differentiation potential (**Figure S1C,D**).

Phenotypical long-term follow-up showed that early hematopoietic recovery, long-term multilineage blood production, as well as reconstitution of the BM stem cell compartment entirely originate from the gene-modified CD34^+^CD45RA^−^CD90^+^ cell subset.

### Multipotent clones contribute to very early neutrophil recovery and persist long-term

To determine the temporal contribution of multipotent CD34^+^CD45RA^−^CD90^+^ clones, PB WBCs, PB subsets, and BM CD34^+^ cells were analyzed by ISA and data aggregated using a new retrospective clonal tracking approach (**Figure S2A**). Briefly, longitudinally tracked clones in WBC samples were retrospectively matched with ISA data from FACS-purified lineages allowing more frequent sampling and reliable capture of clones in the early phases of engraftment even with low WBC counts.

High polyclonality was observed in all animals without any evidence of dominant clones developing (**Figure S2B**). The retrospective lineage tracking approach identified the majority of detected clones in one or more lineages (Z13264: 61.9%, Z14004: 68.5%, Z15086: 70.3%) (**Figure 2A**). Most importantly, we identified thousands of clones contributing to all 5 lineages in Z13264 (1632 clones) and Z14004 (1311 clones). However, the inability to FACS-purify fluorochrome expressing lineage cells for Z15086 led to a significantly lower number of multipotent 5 lineage clones (n=51) despite the high number of total clones detected (**Figure 2A**).

**Figure 2.**
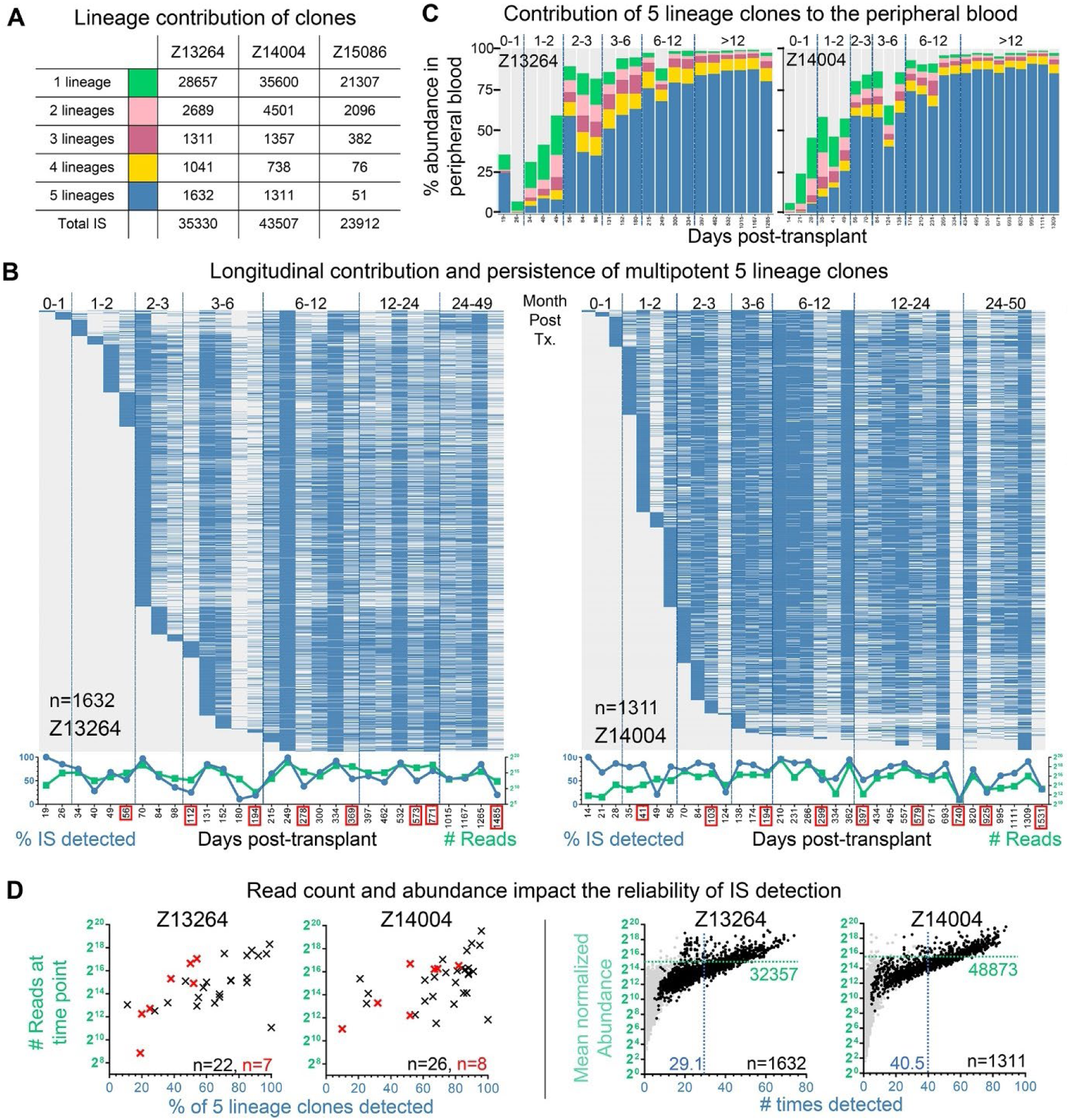
Contribution and persistence of multipotent 5-lineage clones. **(A)** Summary of retrospectively assigned lineage contribution for unique clones found in CD90 animals. Longitudinal detection of multipotent 5-lineages clones. Blue indicates the presence, grey the absence of clones. Graph below heatmap: Frequency of 5-lineage clones detected (blue, left y-axis) and number of reads (green, right y-axis) for each time point. BM time points are highlighted in red. Longitudinal contribution of 5-, 4-, 3-, 2-, and 1-lineage clones to the peripheral blood. Color code as defined in A. Grey portion are clones that were not possible to assign to distinct lineages. **(D)** Impact of data quality and clonal abundance on the reliability of IS detection. Left two graphs: Correlation of the cumulative read count from a single PB (black symbol) or BM (red symbol) time point, with the frequency of multipotent 5-lineage clones detected at the same time point. Right two graphs: Correlation of the mean normalized abundance of 5-lineage clones (black dots), with the number of times the clone was detected across all analyzed samples. Grey background indicates clones associated to other groups. The mean normalized abundance and average number of times detected across all clones are indicated with the green horizontal line and the blue vertical line, respectively.

In contrast to the current literature, multipotent clones were detected in the very first PB WBC samples taken immediately after neutrophil recovery on day 19 and day 14 post-transplant in Z13264 and Z14004, respectively (**Figure 2B**). Multipotent clones found within these early time points contributed long-term and were detected in BM CD34^+^ cells up to 4 years later (**Figure 2B and Figure S2C**).

More than 65% of all 5 lineage clones seen in the entire four years of follow-up contributed to the PB within the first 70 days in Z13264/Z15086 and only 56 days in Z14004 (**Figure 2B, Figure S2D**). Similarly, more than 50% of all mature gene-marked blood cells in the PB were derived from 5 lineage clones by day 70 in Z13264 and Z14004 (**Figure 2C**). The total number of 5 lineage clones detected plateaued at 9 months in Z13264 and Z14004 with no or very few additional multipotent clones detected thereafter.

ISA without purification of gene-marked lineages for Z15086 led to striking differences in the patterns of multipotent HSPC contribution. Clonal tracking data from this animal indicates later as well as lower contribution of multipotent clones to the blood compared to Z13264 and Z14004 (**Figure S2**). While the lower frequency of gene marking did not have any impact on the overall detection of clones in bulk PB WBC samples (**Figure S2A**), the inability to FACS-purify fluorochrome-expressing lineages led to a much lower number of 5 lineage clones detected, highlighting the impact of sampling on the data quality and interpretation.

Longitudinally tracked multipotent clones were not continuously detected throughout the entire follow-up, implying an oscillating contribution and detection over time. Detailed assessment of the data revealed that the reliability of detection was a combined result of the number of samples taken that specific day, the number of sequencing reads, as well as the abundance of the clone in the peripheral blood (**Figure 2B,2D and Figure S2C,S2E**). All three factors were closely interconnected since the number of samples run at a specific day increased the number of sequence reads and consequently enhanced the reliability of detection for low abundance clones.

In summary, we provide evidence that engrafted multipotent HSCs actively contribute from the very first day to hematopoietic recovery without losing their long-term multilineage differentiation potential. Furthermore, multipotent 5-lineage clones are the predominant source of mature gene-marked blood cells as early as 2 months post-transplant. To this end, we predict that the vast majority of 5-lineage clones are continuously contributing to the production of peripheral blood cells.

### Enhanced reproducibility of integration site profiles in CD90 animals

To confirm that enhanced sampling and high density clonal tracking rather than our CD90-mediated gene modification approach results in earlier detection of multipotent HSCs, we comprehensively compared the IS profile of animals transplanted with FACS-purified CD90^+^ cells (CD90 animals) to historic controls receiving CD34^+^ cells (CD34 animals) (**Figure 3A, Figure S3A**). High-level chromosomal mapping of IS demonstrated nearly identical patterns and strong correlation (R^2^=0.607) of transgene localization in both groups (**Figure 3B**). Percentages of 4.8% and 4.2% in-gene IS were found to be associated with oncogenes in CD34 and CD90 animals, respectively. The majority of IS near oncogenes was shared, no correlation was seen in the abundance, and no clonal outgrowth was associated with these IS in between both groups (**Figure 3B, Figure S2B**).

**Figure 3.**
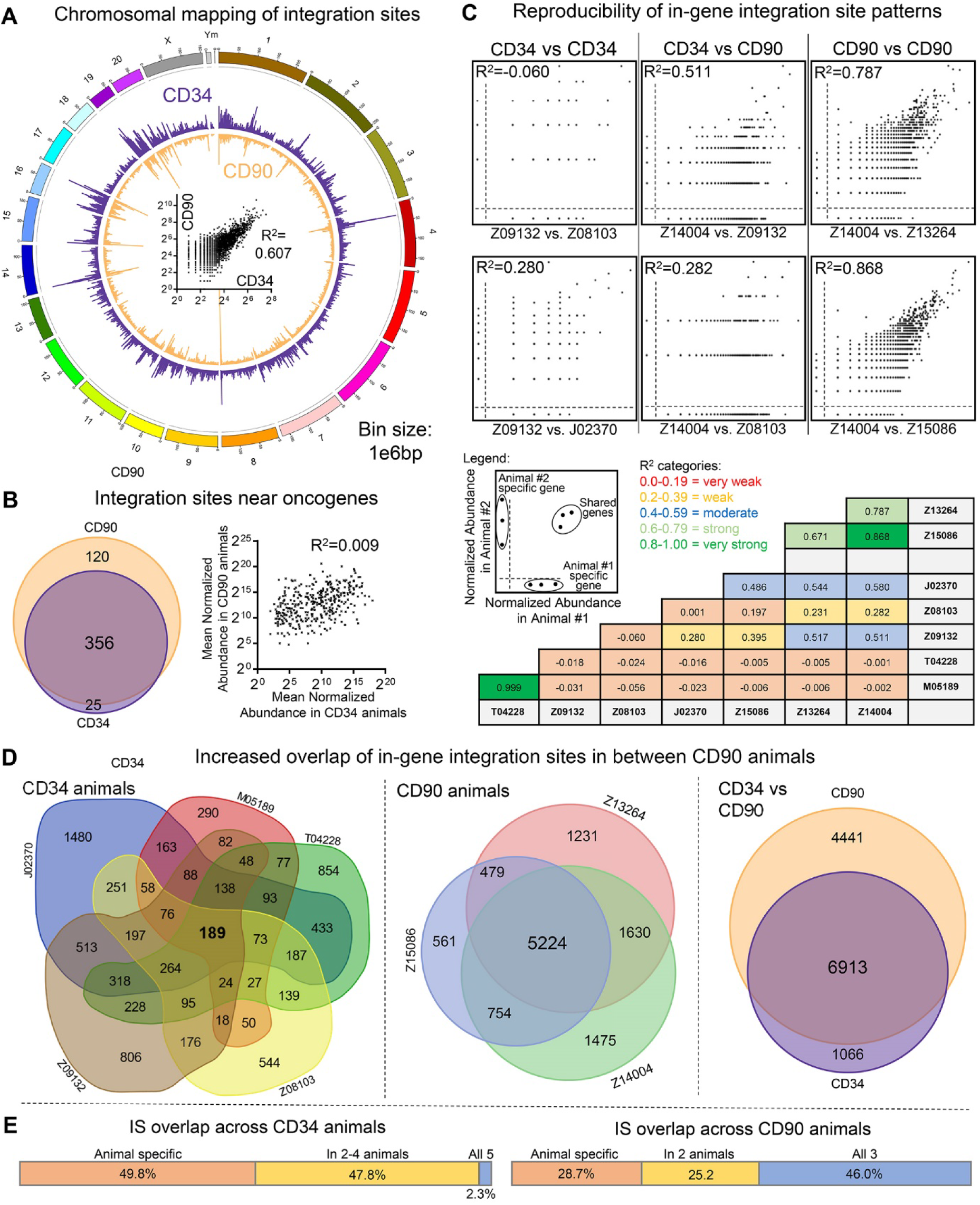
Comparison of integration site profiles in CD34 and CD90 animals. **(A)** Chromosomal mapping of IS distribution in CD34 (purple outside ring) and CD90 (orange inside ring) animals. Bar height indicates the cumulative number of unique IS across all animals within a 1e6 base pair bin. Center graph: correlation of the number of IS within each bin across both groups. **(B)** Left: Comparison of shared and unique IS near oncogenes in between CD34 and CD90 animals. Right: Correlation of the mean normalized abundance of shared IS near oncogenes in between CD34 and CD90 animals. **(C)** Pair-wise comparison of the normalized abundance of in-gene IS. For genes with multiple unique IS, normalized counts were combined. **(D)** Euler and Venn diagrams of in-gene IS overlap in between animals within each group (left and middle) as well as across CD34 and CD90 animals (right). **(E)** Percent of in-gene IS that were animal specific (red), partly shared (orange), and shared in between all animals of one group (blue) for CD34 (left) and CD90 animals (right).

We next determined the reproducibility of IS patterns comparing the data of individual animals within and across both groups. IS associated with the same gene were clustered and the correlation calculated for each pair-wise comparison of animals (**Figure 3C, Figure S3B,C,D**). Correlations in between CD34 animals were generally weak or very weak due to the vast majority of genes not being shared. A weak to moderate correlation was reached when CD34 animals were compared to CD90 animals. A strong to very strong correlation was only seen for the comparison among CD90 animals, highlighting very high reproducibility of in-gene IS patterns (**Figure S2A**). Overlap of in-gene IS across animals was further visualized creating Euler/Venn diagrams (**Figure 3D,E**). Almost 50% of in-gene IS were uniquely found in individual CD34 animals with as little as 2.3% found in all five, whereas 46% of all in-gene IS were shared across CD90 animals with 28.7% found only in individual animals. Finally, the vast majority (86.6%) of IS found in CD34 animals was detected in the CD90 cohort.

High similarity of IS profiles between animals transplanted with genetically modified CD34 and CD90-enriched cells confirmed that historically described lentiviral IS patterns were not altered due to the direct gene-modification of CD34^+^CD45RA^−^CD90^+^ cells. Low density sampling in CD34 animals limited the reproducibility of IS patterns in between animals, whereas high density sampling of CD90 animals significantly enhanced the resolution and reproducibility of IS patterns across animals.

### Detection limit of ISA and low clonal abundance cause multipotent clones to appear lineage restricted

Due to the observed impact of sampling on the reliability of clone tracking, we closely examined all clones in CD90 animals that were found to contribute to only 2, 3 or 4 lineages (**Figure 2A**). Longitudinal tracking demonstrated long-term persistence of many 4- and 3-as well as some 2-lineage clones with similar patterns of contribution as previously described for multipotent 5-lineage clones (**Figure 2B, S2C, S4A**). The reliability of IS detection (number of times seen across all samples) decreased stepwise from multipotent to 1-lineage clones (**Figure 2D, 4A**). None of the 4-, 3-, and 2-lineage clones in Z13264 and Z14004 were highly abundant, and the mean abundance across all clones in each group decreased stepwise from multipotent to 2-lineage clones (**Figure 2D, 4A**). Furthermore, no obvious enrichment for distinct types of lineage-restricted clones was found, indicating that the detection of such clones is more likely a stochastic event (**Figure 4C**). Since none of the most highly abundant and confidently detected clones were found to be lineage restricted (**Figure S4A**), we believe the clones that appear to be lineage restricted are merely undersampled multipotent clones.

**Figure 4.**
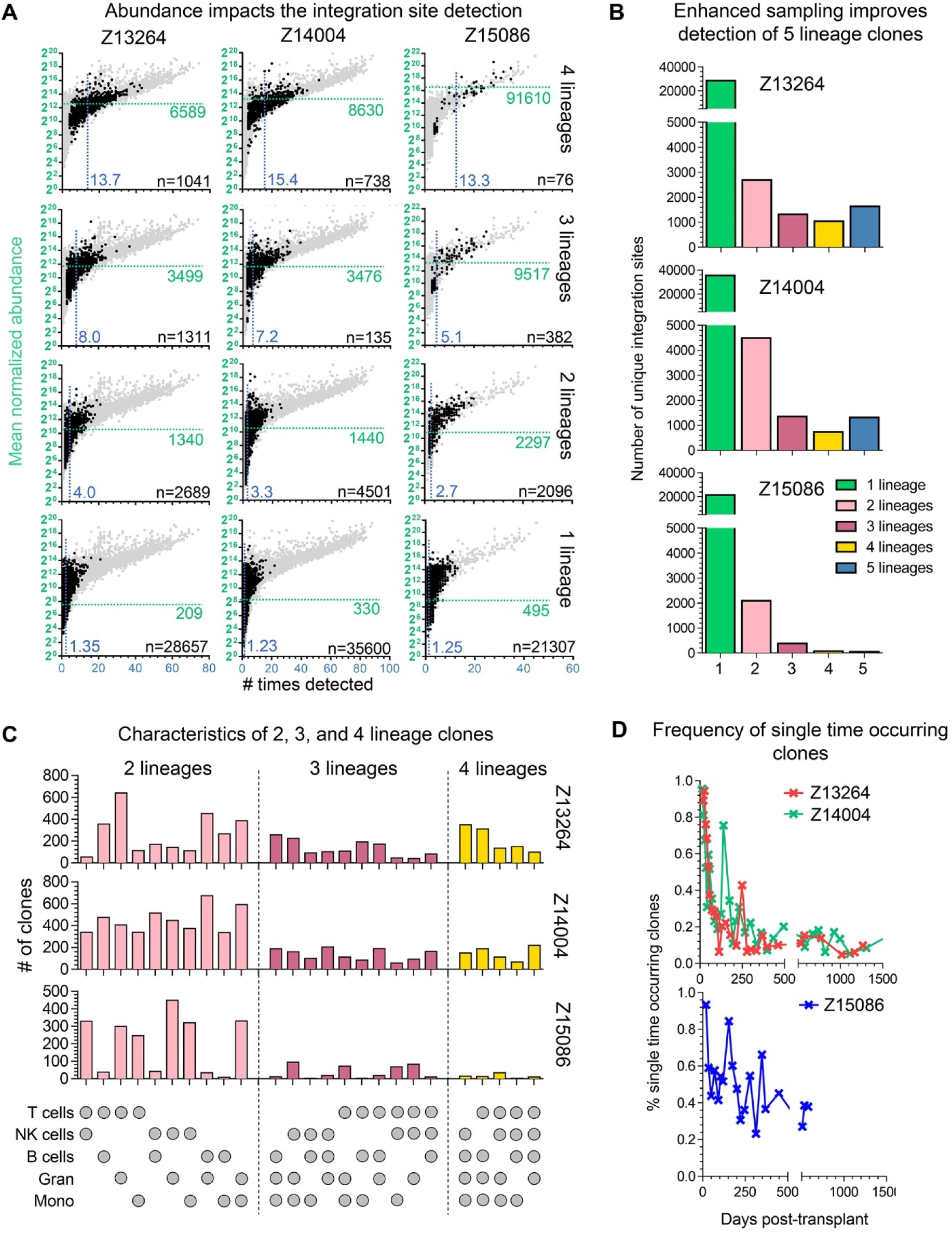
Contribution of lineage-restricted and single-time occurring clones. **(A)** Correlation of the mean normalized abundance of 4-, 3-, 2-, and 1-lineage clones with the number of times the clone was detected across all analyzed samples. Grey background indicates clones associated with other groups. The mean normalized abundance and average number of times detected across all clones are indicated by the green horizontal line and the blue vertical line, respectively. **(B)** Frequency of singly-occurring clones over time. **(C)** Quantification of the different types of 2-, 3-, and 4-lineage clones. **(D)** Longitudinal frequency of single time occurring clones found in the peripheral blood relative to the total number of clones detected at each time point.

The effect of sampling and high density tracking of purified gene-marked cells becomes evident in animal Z15086 where the frequency of gene-modification was below 5%, no FACS-purification of gene-marked lineages was possible, and only four PB subset purifications were performed. As a result, the number of clones to be assigned in Z15086 to a lineage stepwise decreased from 2- to 5-lineage clones (**Figure 2A,4B**). In stark contrast, more multipotent HSC clones than 4- or 3-lineage clones were found in Z13264 and Z14004 (**Figure 2A,4B**). Finally, the detection of 2-, 3-, and 4-lineage clones was affected by the lower data density, suggesting the preferential existence of certain lineage combinations (**Figure 4C**).

We further analyzed clones only found to contribute to a single lineage as well as single-time occurring (SO). The majority of single-lineage clones were found within the first 6-9 months with only very few clones persisting long-term (**Figure S4B**). The majority of 1-lineage clones had a very low abundance, 1.23-1.35 samples on average, and strongly overlapped (Z13264: 54%, Z14004: 57%) with the group of SO clones. SO clones were the predominant source for cells in the PB for the first 50 days after transplant in all three animals (**Figure 2C, S2D**, grey portion). Although we were not able to assign these early SO clones to distinct lineages, flow-cytometric analysis of the PB showed that granulocytes were the predominant cell type during this phase of recovery and, therefore, the only cell fate realized by CD34^+^CD45RA^−^CD90^+^ derived SOs (**Figure 1D**). SO clones were continuously detected at low frequencies (5-10%) in Z13264 as well as Z14004 throughout the entire follow-up, whereas SO frequencies remained at 20 to 40% in animal Z15086, suggesting a sampling related effect (**Figure 4D**).

Based on these data, we hypothesize that all long-term persisting clones detected in 2, 3 or 4 lineages equally originate from multipotent 5-lineage clones, with an abundance close to the detection limit of ISA. More importantly, early hematopoietic recovery is almost entirely driven by a large number of CD34^+^CD45RA^−^CD90^+^ derived SOs.

### Rapid decrease of clonal diversity as a result of stochastic HSC expansion and differentiation

The observed early wave of neutrophilic singlets derived from the HSC-enriched CD34^+^CD45RA^−^CD90^+^ subset (**Figure 4D**), the rapid decrease of clonal diversity in Z13264 and Z14004 (colored lines, **Figure 5A**), and the early contribution of multipotent long-term persisting HSCs during neutrophil recovery do not support the bi-phasic model of recovery. To understand the kinetics of the observed changes in clonal diversity *in vivo*, we simulated the clonal outgrowth and differentiation of CD34^+^CD45RA^−^CD90^+^ cells.

**Figure 5.**
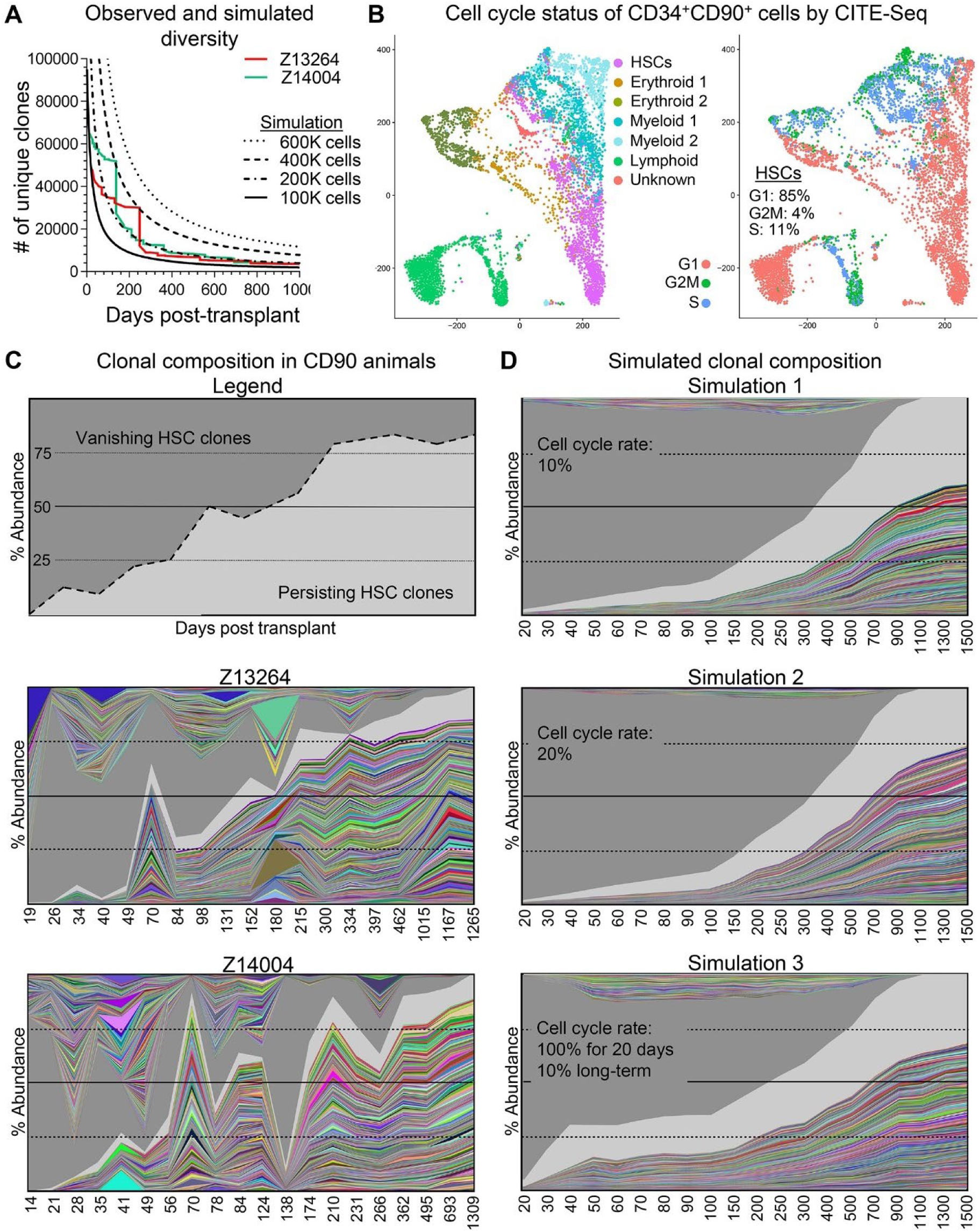
Simulation of a stochastic HSC engraftment. **(A)** Overlay of observed clonal diversity in Z13264 (red line) and Z14004 (green line) with the simulated clonal diversity (black lines). For the simulated diversity different starting cell numbers of engrafted HSCs were used as indicated in the figure legend. **(B)** CITE-Seq of a representative human steady-state bone marrow CD34^+^ population. Transcriptionally-distinct progenitor subsets were color coded as indicated in the figure legend (left graph). Cell cycle status of cells (right graph). **(C)** Longitudinal abundance of short- and long-term persisting clones in Z13264 and Z14004. Clones with greater than 1% are color coded; all other clones with less than 1% contribution are shown in grey shades. **(D)** Simulated longitudinal abundance of short- and long-term clones with variations of the cell cycle rate as indicated in the plot. Color coding as described in C.

Based on our previous findings, we used the following parameters for the simulation: **1)** we assumed all CD34^+^CD45RA^-^CD90^+^ cells to have the same proliferation and differentiation potential at any time; **2)** we assumed that the ratio of self-renewal vs. differentiation is population neutral (50%:50%) in order to maintain the pool of multipotent HSPCs (**Figure S5A**); **3)** we used CITE-Seq (cellular indexing of transcriptomes and epitopes by sequencing) on steady-state bone marrow-derived bulk CD34^+^ cells to determine the cell cycle status of CD34^+^CD90^+^ cells (**Figure 5B, S5B**). CITE-Seq was performed on human CD34^+^ cells due to lack of commercially available NHP cross-reactive CITE-Seq antibodies. On average, 20-25% of CD34^+^CD90^+^ cells were determined to be in a state of proliferation, with 21% in S phase and 4% in G2M. **4)** Finally, we defined the starting cell number for our simulation. Z13264 and Z14004 both received approximately 500-600K gene-modified CD34^+^CD90^+^ cells at the day of infusion^7^.

With 600K gene-modified CD34^+^CD90^+^ cells, the simulated temporal change of clonal diversity was significantly offset from the observed diversity in Z13264 and Z14004 (dotted line, **Figure 5A**). In particular, the number of persisting clones was predicted to be at almost 10,000 clones and, therefore, more than 3-fold higher than the number of persisting HSC clones observed in both animals (**Figure 5A**). We therefore reduced in a stepwise fashion the number of engrafted CD34^+^CD90^+^ cells to 400K, 200K as well as 100K, assuming a portion of HSCs is not homing and engrafting in the bone marrow. The best fit of our model was seen in between 100K and 200K engrafted CD34^+^CD90^+^ cells. Approximately 120K engrafted CD34^+^CD90^+^ cells closely matched the observed number of 3,000 to 4,000 long-term clones *in vivo*.

Next, we tested whether a stochastic engraftment of 120K CD34^+^CD90^+^ cells would actually match the observed contribution of vanishing and persisting HSCs clones contributing to the PB of Z13264 and Z14004 (**Figure 5C**). For this comparison, the hypothetical blood production of vanishing and persisting CD34^+^CD90^+^ clones was simulated (see methods for details). Parameters for this simulation are summarized in the Methods section. With a constant cell cycle rate of 10% for CD34^+^CD90^+^ cells, dominant contribution of persisting HSC clones was significantly delayed to more than 300 days post-transplant in comparison to 50-70 days in Z13264 and Z14004 (Simulation 1, **Figure 5D**). Surprisingly, an increase of the cycling rate to 20% had no significant impact on the clonal composition (Simulation 2). We next tested a two-step model with 100% cycling rate during a 20-day neutrophil recovery phase and 10% cycling thereafter for steady-state hematopoiesis (Simulation 3). The higher cycling rate during recovery did enhance the contribution of persisting HSC clones, whereas dominance for the blood production remained delayed at 200 days post-transplant. We show that the change of clonal diversity seen *in vivo* follows a stochastic pattern, providing evidence that single-lineage neutrophils during the early phase of recovery are HSC-derived. Furthermore, simulating the contribution of HSCs to the peripheral blood, we were able to partly match the detection of vanishing and persisting HSC clones observed by ISA *in vivo*. Our simulations demonstrate that the number of HSC clones rapidly declines over time with only few clones generating bigger long-term persisting pools (**Figure 6**).

**Figure 6.**
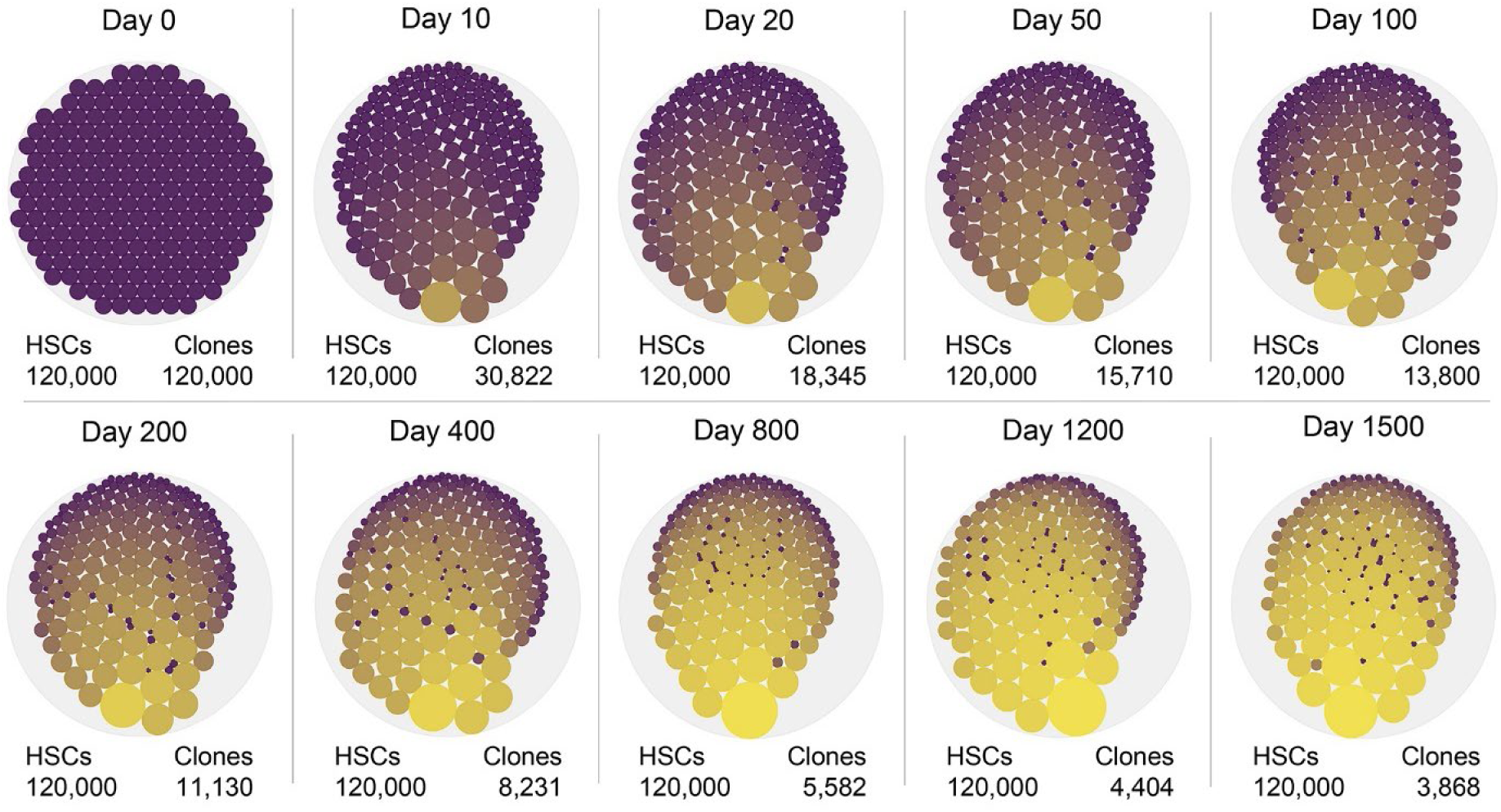
Longitudinal change of clonal diversity and HSC clone size. Visualization of the simulated change of diversity in the bone marrow stem cell compartment. Simulations were performed with 120,000 HSCs at the time of transplant, an initial cell cycle rate of 100% for 20 days, and 10% cycle rate thereafter. The number of clones for each representative time point is shown on the top right. Dot sizes illustrate the clone size.

### Stochastic *in vitro* expansion and differentiation of CD34^+^CD45RA^−^CD90^+^ cells

Our previous simulation uses the tracking of clones in the PB as a surrogate to reverse model a stochastic engraftment of HSCs in the BM. No actual BM samples for ISA on CD34^+^ cells were taken during the first 6 to 8 weeks post-transplant to prevent the depletion of HSC clones and not disrupt the clonal composition before clonal pools are established. To mimic the BM recovery and test whether CD34^+^CD45RA^−^CD90^+^ cells follow our presumed stochastic outgrowth, *in vitro* experiments in colony-forming cell (CFC) assays were carried out.

Initially, we determined the cell cycle rate of CD34^+^CD45RA^−^CD90^+^ cells in CFC assays to adjust our simulation of a stochastic proliferation and change of diversity for CD34^+^CD45RA^−^ CD90^+^ cells in this environment. Similar to our transplant protocol, CD34^+^ cells from two NHPs were enriched (day -1), primed *ex vivo* overnight, modified with CFSE (day 0), and rested overnight. CFSE^+^CD34^+^CD45RA^−^CD90^+^ cells were FACS-sorted on day 1 into CFC medium, harvested every 24 hours for 4 days, and the CFSE intensity was determined flow-cytometrically. On average, the vast majority of CD34^+^CD45RA^−^CD90^+^ cells underwent one division within 24 hours and continued to proliferate at this rate for the consecutive days (**Figure 7A, S6A**).

**Figure 7.**
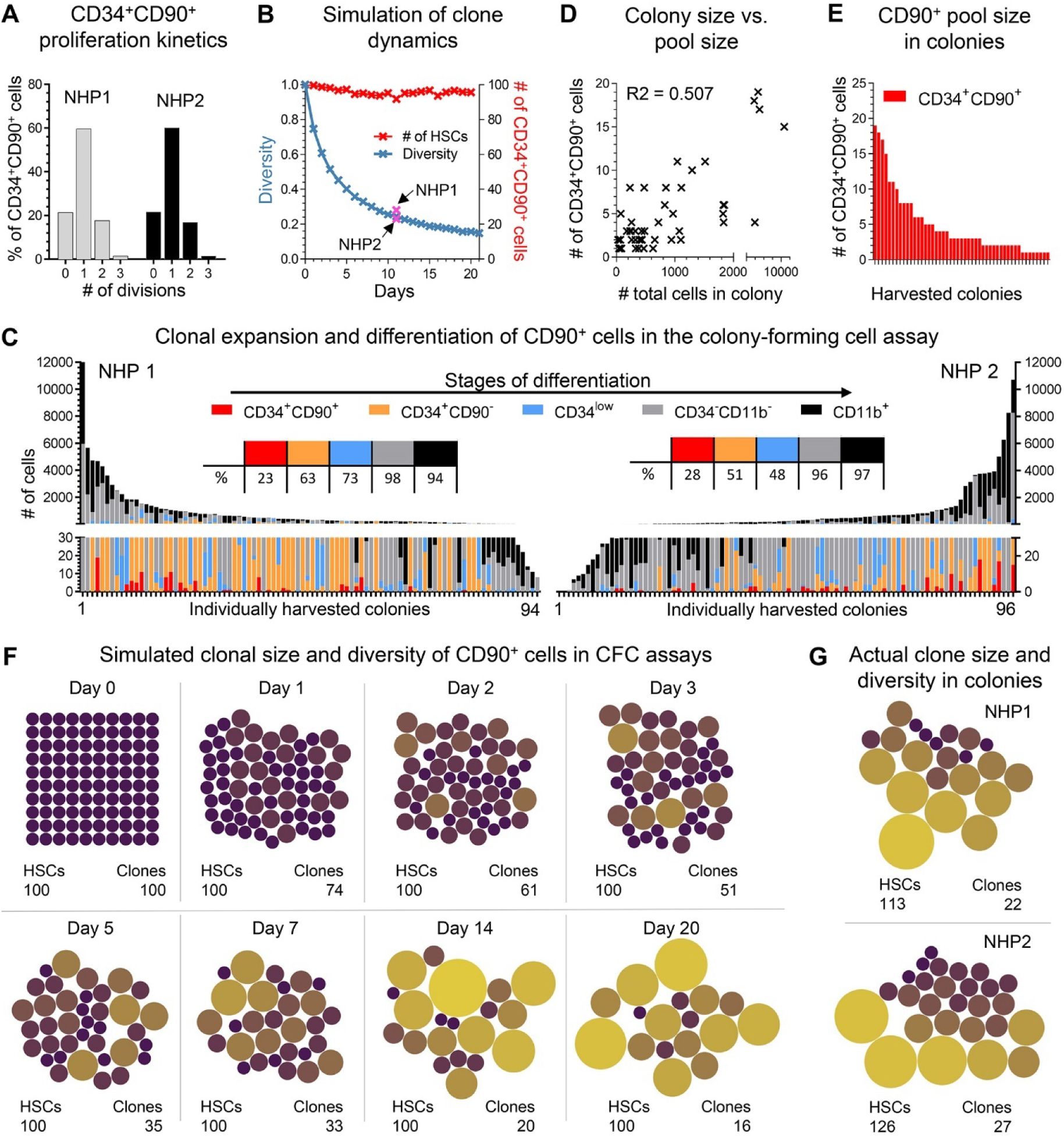
Stochastic outgrowth of CD34^+^CD90^+^ cells in CFC assays. **(A)** Number of divisions of CFSE stained CD34^+^CD90^+^ cells after 24-hour culture in CFC medium. CFSE staining was determined flow cytometrically, and the number of divisions as well as the frequency of divided cells were calculated relative to the parental population. **(B)** Simulation of the change in diversity (blue line) and number of CD34^+^CD90^+^ cells in colonies over time. **(C)** Size and composition of CD34^+^CD90^+^-derived colonies after 11 days of culture. Phenotypes and cell numbers were determined flow-cytometrically. Tables in the center indicate the frequency of colonies containing phenotypically-defined stages of differentiation defined above. **(D)** Correlation of the size of CD34^+^CD90^+^-containing colonies with the actual number of CD34^+^CD90^+^ cells within each colony. **(E)** Number of CD34^+^CD90^+^ cells within colonies. **(F)** Visualization of the change in clonal diversity and CD34^+^CD90^+^ clone size in colonies based on the simulation. **(G)** Actual clonal diversity and CD34^+^CD90^+^ clone size from the CFC assay in C. The dot size is proportional to the number of cells in the colony.

We created a simulation for the outgrowth of CD34^+^CD45RA^−^CD90^+^ cells in CFC assays based on the 24-hour cycle rate. The simulation showed a rapid decrease of diversity within the first few days and loss of more than 80% of the seeded clones by day 15 (**Figure 7B**). The cumulative total number of CD34^+^CD45RA^−^CD90^+^ cells remained the same due to the population neutral assumption of this simulation (**Figure 7B**).

Next, single CD34^+^CD45RA^−^CD90^+^ cells after 48 hours culture *ex vivo* were FACS-sorted into CFC assays, grown for 11 days, individual colonies harvested, and the total number of cells as well as the phenotype of progenies determined (**Figure 7C**). Colonies as large as 12,000 cells were found, and the composition of colonies was a highly diverse set consisting of primitive HSPCs, early progenitors, non-committed cells, and mature myeloid cells (**Figure S6B**). Primitive CD34^+^CD90^+^ HSPCs were found in 23% (NHP1) and 28% (NHP2) of colonies. No obvious clustering was found for colonies containing CD34^+^CD90^+^ cells based on size or composition that would indicate the presence of functionally distinct subsets having different expansion or differentiation potential (**Figure S6C**). Similarly, greater quantities of CD34^+^CD90^+^ cells were not associated with smaller, less-proliferative colonies and correlated with the total size of colonies (**Figure 7D,E**, R^2^=0.507). The cumulative number of CD34^+^CD90^+^ cells across all colonies was above the number of cells seeded (NHP1: 119%, NHP2: 131%), indicating a slight expansion of CD34^+^CD90^+^ cells.

The simulated stochastic reduction of clonal diversity and expansion of some CD34^+^CD90^+^ clones in CFC assays is illustrated as bubble plots in **Figure 7F**. As illustrated in Figure 7B, the number of clones rapidly declines with a small number of individual clones growing into bigger pools. As expected, the simulated diversity closely matched the experimentally determined number of clones as well as the size of CD34^+^CD90^+^ clone pools in CFC assays (**Figure 7G**). Overlaying the observed frequency of colonies containing primitive CD34^+^CD90^+^ HSPCs in Figure 6C with our prediction matched the expected diversity of 24.4% at day 11 (**Figure 7B**).

In summary, the clonal outgrowth and differentiation of CD34^+^CD45RA^−^CD90^+^ cells in CFC assays is a stochastic process. This data supports our previous assumption that the engraftment and contribution of CD34^+^CD45RA^−^CD90^+^ cells *in vivo* in the bone marrow stem cell compartment is similarly a stochastic event rather than a bi-phasic recovery driven by short-term progenitors and long-term HSCs.

## DISCUSSION

Here, we show evidence that hematopoietic reconstitution after myeloablative conditioning and transplantation is primarily driven by multipotent HSCs. Enhanced sampling and increased density of clonal tracking data demonstrate that long-term engrafting HSCs actively contribute during neutrophil recovery and are the predominant source of mature blood cell production as early as 50 days post-transplant. Most importantly, observed changes in the clonal diversity during early recovery suggest a stochastic HSC engraftment and differentiation pattern rather than a bi-phasic reconstitution initially driven by short-term progenitors.

Our high-density ISA data challenges the currently widely accepted two-phase model of hematopoietic reconstitution that assumes contribution of multipotent HSCs earliest at 6-9 months post-transplant^2,3^. Our studies provide evidence that long-term persisting HSCs are actively contributing to neutrophil recovery while maintaining long-term multilineage engraftment potential. The abundance of multipotent clones during neutrophil recovery is very low and, due to the poor sensitivity of ISA, can be easily missed when samples are taken too late or several months apart. In addition, the very rapid decline of diversity during neutrophil recovery with thousands of clones contributing only for a very short time and with low abundance requires a very high sampling density to reliably detect the contribution of multipotent HSCs at these early time points. Such comprehensive sampling would be prohibitive in patients and is only possible in large-animal models such as NHPs. While samples from gene therapy patients were consistently taken throughout the long-term follow-up, early sampling was limited to monthly draws with the first samples available for analysis 4-6 weeks post-transplant^2^. NHP studies similarly focused on the long-term tracing of clones with limited data density, specifically in the first 3 months post-transplant, likely missing the vast majority of HSC-derived clones contributing during early recovery^3,4^. A significant benefit of clonal tracking in large animals such as NHPs is the ability to purify gene-modified cells based on their fluorochrome expression. The purification of gene-modified WBCs during early time points especially improved the quality of ISA data and increased the number of multipotent 5-lineage clones we were able to detect. The inability to purify gene-modified lineages from Z15086 due to the animal’s low weight, the allowed blood volume to collect, as well as the low gene-marking levels led to a significantly lower detection of multipotent HSCs in comparison to animals Z13264 and Z14004, illustrating the importance of sample purity.

Although we enhanced the resolution of our ISA dataset, we are still experiencing sample-to-sample variability. High density sampling as well as the abundance of clones had a significant impact on the reliability of detection. Individual time points with poor sample quality or low number of sequencing reads only detected highly abundant clones and introduced a seemingly oscillating contribution or even lineage bias of multipotent HSCs. While other groups have reported the presence of lymphoid- and myeloid-biased HSC clones ^2,5^, we provide evidence that lineage skewing may be a result of the sensitivity of ISA. Although Six et al. addressed this issue by rerunning the same time point multiple times and increasing the sensitivity of ISA, the limited material from patients only contains a fraction of contributing clones and does not represent the entire repertoire of clones present at a given time point. Furthermore, the outgrowth of individual HSC clones forming large pools of identical siblings makes it difficult to decipher whether a single HSC from this pool may be biased in the lineage output. In order to reliably address this question, new tracing methods that capture the number of HSC divisions and introduce a novel fingerprint into identical HSC daughter cells will be needed. Transposase-based systems as used by Sun et al. that are combined with a cell-cycle activated promoter may enable the tracing of otherwise identical HSCs^8^.

The general assumption is that HSC-enriched CD34 subsets such as the CD34^+^CD45RA^−^CD90^+^ phenotype or others contain a mix of different cell entities^7,9,10^ similar to short-term and long-term HSCs previously reported in mice^11^. However, scRNAseq did not reveal any transcriptionally distinct subsets within such phenotypically defined, HSC-enriched CD34 populations that would support the coexistence of two or more functionally distinct subsets with short- and long-term engraftment features^12,13^. The initial rapid decline in clonal diversity in combination with the long-term persistence of only a couple thousand clones rather indicates a stochastic reconstitution driven by a single cell entity. Simulations of a stochastic recovery did match the observed clonal diversity *in vivo* and even predicted the clonal outgrowth of CD34^+^CD45RA^−^CD90^+^ cells in CFC assays *ex vivo*. Our data therefore implies that all CD34^+^CD45RA^−^CD90^+^ cells engrafting and homing into the BM are identical in their potential and that the decision to either contribute early to neutrophil recovery as a single time occurring clone or to expand and persist building up a pool of identical HSC clones is intrinsically controlled. However, our simulations suggest that only 120K out of 600K transplanted gene-modified CD34^+^CD45RA^−^CD90^+^ cells contributed to hematopoietic recovery. It remains unknown if the missing 70-80% failed to home to the BM, differentiated before neutrophil recovery, or were simply missed due to their low abundance during the early days of recovery. Future research and technological advances in clone tracking and scRNAseq technologies may enable us to determine the fate and potentially identify subsets within CD34^+^CD45RA^−^CD90^+^ cells that may be associated with these “missing” clones.

Our simulations are based on a limited number of parameters that do not include some factors that have been previously applied to other HSC engraftment models. Factors that have been proposed to have an impact on the proliferation of HSCs include positive/negative feedback mechanisms of the tissue and mature blood cells, differences in the symmetric/asymmetric divison rate, as well as the capacity limit of the BM niche^14^. Since changes in the ratio of symmetric vs. asymmetric division are frequently associated with a loss or gain of stem cells either leading to exhaustion or an excessive and potentially cancerogenic outgrowth^15-19^, we consider the cell cycle rate to be the most important parameter. Introduction of environmental regulators as well as the BM capacity feedback into our simulation are expected to modify the cell cycle rate and, presumably, the speed of change in clonal diversity specifically in the early phase of engraftment. We currently assume that all CD34^+^CD45RA^−^ CD90^+^ cells cycle for 20 days in our model, but *in vivo* feedback mechanisms may gradually slow down HSCs until steady-state hematopoiesis is reached. The temporal change of the cell cycle rate for the CD34^+^CD45RA^−^CD90^+^ subsets after myeloablation and transplant is currently unknown and will require more research in order to refine our simulation and further improve upon our current predictions. Furthermore, we would expect to see a light expansion of HSCs over time to fill in for lost/apoptotic cells or fill empty niches in the bone marrow. First attempts changing our initially assumed ratio of expansion vs. differentiation partly helped to match the clonal dynamics seen in the peripheral blood.

Various models of HSC contribution during steady-state hematopoiesis and recovery after transplantation have been proposed. The common assumption has long been that HSCs are quiescent during steady-state hematopoiesis and recovery after transplantation, rarely contribute to everyday blood production, and have a very limited number of cell cycles before they imminently differentiate^8,20^. This led to the assumption that steady-state hematopoiesis is driven by multipotent progenitors (MPPs) to prevent exhaustion and consumption of HSCs. However, a variety of recent studies in mice actually determined robust and continuous everyday contribution of HSCs to the blood, challenging the idea of a MPP-driven steady-state hematopoiesis^21-24^. Contribution of HSCs during steady-state hematopoiesis was further found in baboons, estimating 56-84% of HSCs to cycle at least once a year^25^. Similarly, contribution of HSCs during hematopoietic recovery after transplant in cats had been proposed more than two decades ago by Abkowitz et al.^26^. Models predict that the engraftment, contribution, and differentiation of HSCs is likely a stochastic event in mice^27,28^, cats^26^, primates^29^, as well as humans^14,30^. Most modelling approaches use observed patterns in the peripheral blood (cell composition, marking efficiency, label retention) as a surrogate to estimate the contribution of HSCs. Here, we demonstrate for the first time that the presumed stochastic engraftment of HSCs *in vivo* can also be applied to single-cell cultures *ex vivo*. The stochastic decision of self-renewal vs differentiation seems to be deeply embedded in HSCs regardless of the environment, external factors, or manipulation.

A stochastic engraftment of HSCs would further impact the interpretation of single-cell transplantation studies with the goal to identify the phenotype of human and murine HSCs^10,31,32^. About 40-46% of single murine and 28% of human phenotypically defined, HSC-enriched cell subsets did demonstrate long-term multilineage engraftment in mice, leading the authors to assume an enrichment rather than isolation of HSCs. However, assuming a stochastic likelihood of long-term engraftment, no more than 30-40% of transplanted cells would be expected to successfully reconstitute mice. Consequently, the previously described phenotypes are presumably much more highly enriched for HSCs than currently assumed, and the chase for additional cell surface markers to identify the supposed subset and yield an even higher purity of HSCs may not succeed.

Our assumption that hematopoietic recovery is primarily driven by HSCs has major implications for the field of allogeneic as well as autologous transplants of HSCs. While we have shown a correlation of neutrophil and platelet recovery with the number of transplanted CD34^+^CD45RA^−^CD90^+^ cells in autologous transplants in NHPs, correlation of the number of human cord blood-derived and expanded CD34^+^CD90^+^ cells with the hematopoietic recovery in children has been shown. Despite the predictive feature of CD34^+^CD90^+^ cell quantification in infusion products and the known variability of this subset in CD34 populations, the current quality control standard of stem cell grafts still relies on the determination of total CD34^+^ cells. Quantification of CD34^+^CD90^+^ cells instead would enhance the quality control of infusion products.

In summary, high density clonal tracking in NHPs revealed an unexpected early contribution of HSCs after myeloablative conditioning and transplantation challenging the current model of a biphasic hematopoietic recovery. This surprisingly robust and continuous contribution of HSCs suggests that hematopoietic reconstitution is primarily mediated by multipotent CD34^+^CD45RA^−^CD90^+^ HSCs. These findings have major implications for HSC transplantation and particularly for HSC gene therapy. The ability to directly modify CD34^+^CD90^+^ cells would substantially reduce the use of expensive reagents needed for the genetic engineering of cells. We here confirmed in the NHP that the direct modification of this HSC-enriched subset *ex vivo* has no negative impact on the overall lentiviral integration site pattern nor increases the risk for potentially oncogen near events, rendering this HSC-targeted gene therapy approach efficient and safe for future clinical application. Similarly, lentiviral transduction of FACS-purified CD34^+^CD45RA^−^CD90^+^ human cells did significantly enhance the gene-modification efficiency and long-term multilineage engraftment in the mouse xenograft model^33^. Most importantly, this phenotype combining CD34 and CD90 is highly unique with currently no other known cell type co-expressing both markers providing a target for highly-specific *in vivo* HSC gene therapy approaches.

## MATERIALS AND METHODS

### Autologous nonhuman primate transplants and *ex vivo* engineering of HSPCs

Autologous nonhuman primate transplants, priming (mobilization), collection of cells and genetic engineering were conducted as previously reported ^7,34^. All experimental procedures performed were reviewed and approved by the Institutional Animal Care and Use Committee of the Fred Hutchinson Cancer Research Center and University of Washington (Protocol #3235-01). This study was carried out in strict accordance with the recommendations in the Guide for the Care and Use of Laboratory Animals of the National Institutes of Health (“The Guide”) and monkeys were randomly assigned to the study. Detailed protocols for the performed nonhuman primate transplants including i) the *ex vivo* engineering of HSPCs, ii) animal housing and care, as well as iii) preconditioning and supportive care were previously described ^7^.

### Lentiviral Vectors

The vector used in this study (pRSCSFFV.P140K.PGK.eGFP/mCherry/mCerulean-sW) is a SIN LV vector produced with a third-generation split packaging system and pseudo-typed by the vesicular stomatitis virus G protein (VSVG). The vector for this study was produced by our institutional Vector Production Core.

### Bone marrow harvest and CD34+ enrichment

Bone marrow was harvested and CD34+ cells enriched for flow-cytometry and FACS-sorting as previously described ^34^. Briefly, before enrichment red blood cells were lysed in ammonium chloride lysis buffer, and white blood cells were incubated for 20 minutes with the 12.8 immunoglobulin-M anti-CD34 antibody, then washed and incubated for another 20 minutes with magnetic-activated cell-sorting anti-immunoglobulin-M microbeads (Miltenyi Biotech, Bergisch Gladbach, Germany). The cell suspension was run through magnetic columns enriching for CD34+ cell fractions with a purity of 60-80% confirmed by flow cytometry.

### Flow Cytometry Analysis and FACS

Fluorochrome-conjugated antibodies used for flow cytometric analysis and FACS are listed in **Table S1**. Dead cells and debris were excluded via FSC/SSC gating. Flow cytometric analysis was performed on an LSR IIu (BD, Franklin Lakes, NJ), LSRFortessa X50 (BD), FACSymphony A5 (BD), FACSAria IIu (BD), and Symphony S6 (BD). Cells for *in vitro* assays and integration site analysis were sorted using a FACSAria IIu (BD) and Symphony S6 (BD) cell sorter. Post-sort purity was assessed on the FACSAria IIu and Symphony S6 cell sorter reanalyzing at least 500 cells for each sample. Data was acquired using FACSDiva™ Version 6.1.3 and newer (BD). Data analysis was performed using FlowJo Version 8 and higher (BD).

### Colony-Forming Cell (CFC) Assay

For CFC assays, 800-1,200 sort-purified CD34^+^ cells and CD34-subpopulations were seeded into 3.5 ml ColonyGEL 1402 (ReachBio, Seattle, WA). Colonies were scored after 12 to 14 days, discriminating colony forming unit-(CFU-) granulocyte (CFU-G), macrophage (CFU-M), granulocyte-macrophage (CFU-GM) and burst forming unit-erythrocyte (BFU-E). Colonies consisting of erythroid and myeloid cells were scored as CFU-MIX.

### CFSE assay

CD34^+^ cells were stained with CellTrace™ CFSE according to the manufacturer`s recommendation (Thermo Fisher, Waltham, MA, USA). Briefly, CD34^+^ cells at 10e^6^ per ml were incubated with 1ul of CellTrace™ CFSE stock solution for 20 minutes at 37C, washed with PBS, and cultured in StemSpan (STEMCELL Technologies, Vancouver, Canada) medium supplemented with penicillin-streptomycin (PS) (100 U/ml) (Gibco by Life Technologies) and 100 ng/ml of each stem cell factor (PeproTech, Cranbury, NJ, USA), TPO (thrombopoietin; PeproTech), and FLT3-L (Fms-related tyrosine kinase 3 ligand; Miltenyi Biotec). Cells were cultured at 37°C, 85% relative humidity, and 5% CO2.

### Insertion site analysis

#### Sample processing and sequencing

As described in Adair et al. ^6^, processing of gDNA (genomic deoxyribonucleic acid) to amplify integration loci included either LAM (linear-amplification mediated)-PCR (polymerase chain reaction) ^35^ (animal T04228) or MGS (modified genomic sequencing)-PCR ^36^ methods (animals M05189, J02370, Z08103, Z09132, Z13264, Z14004, and Z15086). A variety of next generation sequencing platforms were used depending on date of transplant and sample collection. Platforms included single end 454 GS FLX Titanium (Roche), single end Ion Torrent PGM (Personal Genome Machine), and paired end Illumina Miseq.

#### Data processing

First, forward and reverse reads were stitched using PEAR ^37^ with the -q 30 option to trim sequence reads after two bases with a quality score below 30 were observed. Stitched FASTQ files and raw FASTA files for all sequencing data were filtered using a custom C++ program. Each read was compared to the reference provirus LTR sequence. Reads with <90% match to the LTR sequence were discarded. The LTR sequence was trimmed off of the remaining reads. Reads were then compared with vector sequence (as opposed to genomic insertion sequences). Reads with ≥80% match to the vector sequence were discarded. The remaining reads were output in FASTA format for alignment. Identified genomic fragments were aligned to the rhesus macaque (rhemac8) genome (GenBank: 2701138) to determine the chromosome, locus, and orientation of integration (e.g., Chr14_8020175_+), as the pigtail macaque genome has not been sequenced. rheMac8 was downloaded from the UCSC (University of California Santa Cruz) genome browser (http://genome.ucsc.edu/).

Filtered and trimmed sequence reads were aligned to the reference genome using BLAT (BLAST-like alignment tool) ^38^ with the following options:

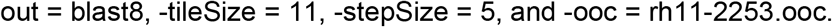

The rh11-2253.occ file contains a list of 11-mer occurring at least 2,253 times in the genome to be masked by BLAT and was generated using the following command:

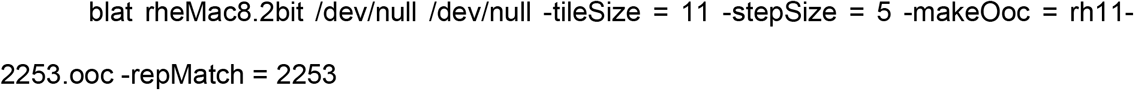

as recommended by UCSC (http://genome.ucsc.edu/goldenpath/help/blatSpec.html and http://genome.ucsc.edu/FAQ/FAQblat.html#blat6). Resulting blast8 files were parsed using a custom python script. Any read with a BLAT alignment length less than 30 or alignment start greater than 10 was discarded. The top scoring alignment and all alignments within 95% of the top score were saved for each read. The ratio of the best alignment to the second-best alignment is the degree with which the insert can be mapped to one location in the genome (multi-align-ratio). Starting with the highest count reads, reads with matching alignments were combined. Reads with multiple possible alignments were not discarded at this point but were grouped together with other reads with the same alignment(s). For each group of reads with matching alignments, the original FASTA sequence files were read, and then the sequences were aligned with Clustal Omega ^39,40^. This alignment was used to build a single consensus sequence for the alignment group. The consensus sequence was then used to search the pool of all sequences that could not be aligned by BLAT, and any sequence with ≥90% identity was merged into the group. Starting with the highest count groups and comparing them with the lowest count groups, groups with ≥90% sequence similarity were merged. Finally, when comparing all sequence files for one test subject, all groups with exact alignment matches were merged into one clone ID. Clone IDs with exact consensus sequence matches were also merged. Non-uniquely aligned groups (multi-align-ratio ≥ 0.9) that had ≥90% similar reference sequences were also merged into a single clone ID. The data was visualized using the R package ggplot26 and custom R scripts.

#### Circos plots

To generate Circos plots (http://circos.ca/)^41^, animals were grouped into two subsets – CD34 and CD90. A list of unique clones and their integration site was generated for each of the animals in the two groups using a custom script in C++. The integration site consists of the chromosome number and the chromosomal location where the vector is integrated (e.g., Chr14_8020175_+). Clones that could not be mapped to specific chromosomal locations (chrUn) were filtered out. All the chromosomes in the rhesus macaque genome were divided into 1,000,000 base pair bins. Integration sites were then mapped to the specific chromosomal location. An input file (.txt) consisting of the chromosome number, location, and the counts of integration sites in that bin (e.g., rm14 8,000,001-9,000,000 18) was generated for each of the CD34^+^ and CD90^+^ groups by combining the data for all the animals within the group. A Circos plot in the form of a histogram indicating the integration sites for the CD34^+^ and CD90^+^ animals along the rhesus macaque genome was generated using the command line version (v.0.69-9).

#### Oncogene mapping

A list of unique clones and their integration sites (e.g., Chr14_8020175_+) was generated for each animal in the CD34^+^ and CD90^+^ groups using a custom script in C++. The Macaca mulatta Mmul_8.0.1.97 GTF file was downloaded from Ensembl (https://www.ensembl.org). The clones were mapped to the reference GTF file using a custom python script to determine if the integration site was in a gene or not. To determine if these genes were oncogenic or not, a database of genes that contain mutations known to be involved in cancer was downloaded from Cancer Gene Census (https://cancer.sanger.ac.uk/census) ^42^.

The genes in the database were classified into two groups – Tier 1 and Tier 2. Tier 1 genes have documented evidence of mutations and activity relevant to cancer while Tier 2 genes are known to have a role in cancer but with less available evidence. The genes associated with the clones were parsed through the database and were classified into Tier 1 or Tier 2 oncogenic groups.

### Cellular indexing of transcriptomes and epitopes by sequencing (CITESeq)

#### Sample preparation

Steady-state BM-derived CD34^+^ cells were stained with TotalSeq™-A antibodies listed in **Table S2** following the manufacturer`s recommendations (BioLegend, San Diego, CA, USA). Briefly, 1e6 CD34^+^ cells in 100ul staining buffer (BioLegend) were blocked with 5ul Human TruStain FcX™ Blocking Reagent for 10 minutes at 4°C (BioLegend). 1ug of each TotalSeq™-A antibody was added to the cell suspension, incubated for 30 minutes at 4°C, and washed 3 times with staining buffer at 800xg for 5 minutes. Antibody stained CD34^+^ cells for single-cell RNA sequencing were processed using the Chromium Single Cell 3’ (v3) platform from 10X Genomics (Pleasanton, CA). Separation of single cells, library preparation, and RNA extraction were performed in accordance with the 10X Chromium Single Cell Gene Expression Solution protocol. To obtain TotalSeq libraries, custom ADT (Antibody Derived Tag) primers (BioLegend) were added to the cDNA amplification reaction. After cDNA amplification ADT-derived cDNA and mRNA-derived cDNA was separated using SPRI select as recommended in the 10X protocol. The cDNA in the supernatant of this reaction was used to generate ADT libraries, whereas the pelleted cDNA was carried forward according to the manufacturer’s recommendation.

#### mRNA Alignment and Counting

The 10X Genomics Cell Ranger software suite was used to convert the raw sequence reads into single-cell gene expression counts. The “cellranger count” command with default options was run for alignment, filtering, cell barcode counting, and UMI counting. cDNA was aligned to human reference genome (hg38) using the STAR aligner (v.2.6.1). UMIs were filtered for a minimum of Qual = 10. Reads were marked as PCR duplicates if two or more read pairs shared the same cell barcode, UMI, and gene ID. Valid cell barcodes were determined based on the final UMI distributions. Valid cell barcodes with a valid UMI mapped to exons (Ensembl GTF GRCh38) were used to generate the final cell barcode matrix.

#### Quantification of ADTs

Antibody-derived tags (ADT) counts were generated from the ADT fastq files using a custom python script, pycite.

#### Dimensional Reduction and Clustering

The single cell data analysis was performed using Seurat ^43^, an R toolkit for single cell genomic data. The CD34^+^ bone marrow RNA counts matrix, and the ADT counts matrix were loaded in RStudio. A Seurat object was created using the RNA data initially. The pre-processing encompassed the selection and filtration of cells based on the QC metrics. Cells with unique feature counts less than 200 and greater than 9000 and mitochondrial counts greater than 20% were filtered out. The ADT counts were added to the object by creating a new assay. The data was normalized using centered log ratio transformation (CLR) and scaled on the ADT assay. The assay of the object was then set to RNA and the data was normalized and scaled again. PCA was run on the highly variable genes to compute linear dimensional reduction. The cells were clustered using a shared nearest neighbor (SNN) modularity optimization-based clustering algorithm with a resolution of 1. Uniform Manifold Approximation and Projection (UMAP) was used to visualize the gene clusters of the object.

#### Cell-Cycle Scoring and Regression

To reduce the effects of cell cycle heterogeneity in the data, cell cycle phase scores were calculated based on markers from ^44^ and were loaded with Seurat. The markers were divided into groups based on G2/M phase and S phase. Each cell in the object was assigned a score, based on the G2/M and S phase markers and the difference between the S scores and the G2/M scores was calculated for the object. The object was scaled again and the difference between the S and G2/M phase scores was regressed out. PCA and UMAP were run on the data, and it was clustered again with a resolution of 1.

#### Cell type module

To determine the cell type of the cells in the data, a feature list was prepared with 3-4 specific gene markers for Stem, Erythroid, Myeloid and Lymphoid cell types. A custom R script was used to assign a cell type to the cells based on the expression of the gene markers.

#### SPRING plots

A custom script was used to calculate the force-directed shared-nearest neighbors ^45^ and a dimensional reduction plot (DimPlot) was used to plot the clusters in the form of force-directed shared-nearest neighbors. A DimPlot was used to visualize the clusters grouped by their cell type and to visualize the cells grouped by their cell cycle phases (G1, S and G2/M). Feature Plots were made to visualize the expression of the AVP and TYMS genes and the CD90 ADT expression. To calculate the percentage of stem cells that are cycling, the cells that were assigned ‘Stem’ cell type were selected. The cell cycle identity (G1, G2/M and S) for those cells was determined and cell cycle rate was calculated.

To visualize the expression of CD90+ ADTs, a histogram of log2 CD90 ADT expression counts in the stem cells was plotted.

#### Next-Generation Sequencing for CITEseq

Sequencing of single cell RNAseq samples was performed using an Illumina HiSeq 2500 in rapid mode employing 26 base read length for read 1 (10x barcode and 10bp Unique Molecular Index (UMI)) and 98 base read length for read 2 (cDNA sequence). Image analysis and base calling was performed using Illumina’s Real Time Analysis (v1.18) software, followed by ‘demultiplexing’ of indexed reads and generation of fastq files, using Illumina’s bcl2fastq Conversion Software (v1.8.4).

### Statistics

Statistical analysis of data was performed using GraphPad Prism Version 5. Significance analyses were performed with the unpaired, two-sided Student’s t-test (*: p < 0.05; **: p < 0.01; ***: p <0.001).

### Software and Packages

FlowJo v.10.2 and higher https://www.flowjo.com

10X Genomics Cell Ranger software suite - v2.0.0 - https://support.10xgenomics.com/single-cell-gene-expression/software/overview/welcome

STAR aligner v.2.6.1 - https://github.com/alexdobin/STAR

Seurat – v 4.0.2 - https://satijalab.org/seurat/

## DATA AND MATERIALS AVAILABILITY

Custom scripts as well as ISA and CITESeq data will be available upon publication or request.

## ACKNOWLEDGEMENTS

We thank Helen Crawford for help in preparing this manuscript and figures.

## AUTHOR CONTRIBUTIONS

SR and HPK designed the study. ME and DP performed ISA data analysis. ME performed modelling. SR and AMP performed CITESeq. DP and ME analyzed CITESeq data. AMP, RM and MC performed longitudinal follow-up, CD34 enrichments, and FACS-sorts. SR, DP and ME generated the figures. HPK funded the study. SR, ME, DP and HPK wrote the manuscript. All authors reviewed and edited the final manuscript.

