## Supplementary material for "Hematopoietic recovery after transplantation is primarily derived from the stochastic contribution of hematopoietic stem cells": Figure S

*both authors contribute equally


### LIST OF SUPPLEMENTAL ITEMS

1. **Table S1.** Antibodies for flow cytometry.
2. **Table S2.** Antibodies for CITEseq.
3. **Figure S1.** Recovery and gene marking of the PB and BM stem cell compartment.
4. **Figure S2.** CD90 animal characteristics, polyclonality in the PB, and clone tracking in animal Z15086.
5. **Figure S3.** Correlation of in-gene integration sites between CD34 and CD90 animals.
6. **Figure S4.** Contribution and persistence of 4-, 3-, 2-, and 1-lineage clones.
7. **Figure S5.** Determination of parameters for the simulation of a stochastic HSC engraftment.
8. **Figure S6.** Proliferation and differentiation of $CD34^+CD90^+$ cells in CFC assays.

### SUPPLEMENTAL TABLES

**Table S1** Antibodies for flow-cytometry.

| Epitope | Clone | Company |
|---|---|---|
| CD3 | SP34-2 | BD Biosciences |
| CD4 | L200 | BD Bioscience |
| CD8a | RPA-T8 | BD Bioscience |
| CD11b | ICRF44 | BioLegend |
| CD14 | M5E2 | BD Biosciences |
| CD16 | 3G8 | BD Biosciences |
| CD20 | 2H7 | BD Biosciences |
| CD34 | 563 | BD Biosciences |
| CD45 | D058-1283 | BD Biosciences |
| CD45RA | 5H9 | BD Biosciences |
| CD90 | 5E10 | BD Biosciences |

**Table S2** Antibodies for CITEseq.

| Epitope | Clone | Company | TotalSeqA barcode | Barcode sequence |
|---|---|---|---|---|
| CD34 | 563 | BioLegend | 54 | GCAGAAATCTCCCTT |
| CD90 | 5E10 | BioLegend | 60 | GCATTGTACGATTCA |

Figure S1

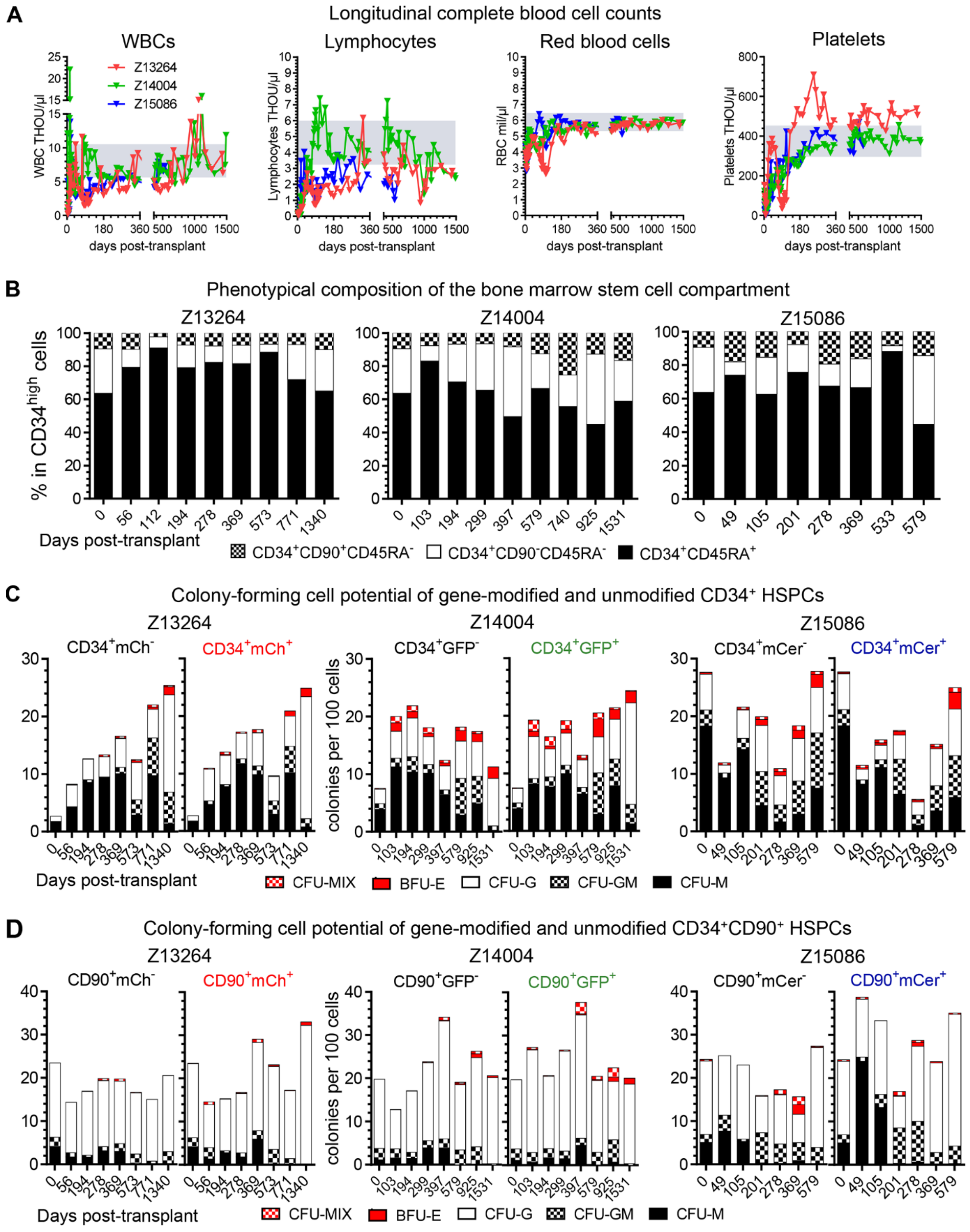


**Figure S1. Recovery and gene marking of the PB and BM stem cell compartment. (A)** Longitudinal complete blood counts (CBCs) measuring white blood cell (WBC) counts, lymphocytes, red blood cells (RBCs), and platelets. Grey boxes highlight the normal range: 5.9–10.4 thousand WBCs per µl, 38-60 thousand lymphocytes per µl, 5.7–6.4 million RBCs per µl, 300–450 thousand platelets per µl. (**B**) Longitudinal flow-cytometric quantification of hematopoietic stem and progenitor cell (HSPC) subsets in the BM. (**C-D**) Colony-forming cell (CFC) potential of unmodified ($GFP^{–}$, $mCh^{–}$, or $mCer^{–}$) and gene-modified ($GFP^{+}$, $mCh^{+}$, or $mCer^{+}$) (B) $CD34^{+}$ as well as (C) $CD34^{+}CD90^{+}$ HSPCs. Unmodified and gene-modified subsets were sort-purified into CFC assays, assays incubated for 10-14 days, and myeloid, erythroid, as well as erythro-myeloid colonies quantified. *Abbreviations:* BFU-E = Burst forming unit-erythrocyte; CFU = Colony-forming unit; CFU-G = granulocyte colony; CFU-GM = granulocyte-monocyte/macrophage colony; CFU-M = monocyte/macrophage colony; CFU-MIX = erythro-myeloid colony; GFP = Green fluorescent protein; mCer = mCerulean; mCh = mCherry.

Figure S2

**A** Integration site data summary

| | Z13264 | Z14004 | Z15086 |
|---|---|---|---|
| % Gene Marking | 25-30% | 30-35% | 1-3% |
| Follow-up [days] | 1485 | 1513 | 579 |
| # Samples | 82 | 104 | 57 |
| # Total IS | 57,027 | 63,539 | 34,009 |
| # IS in Genes | 37,551 | 42,017 | 22,628 |
| # Genes with IS | 8,564 | 9,083 | 7,018 |
| # of Sequencing Reads | 1,908,918 | 3,020,158 | 655,479 |

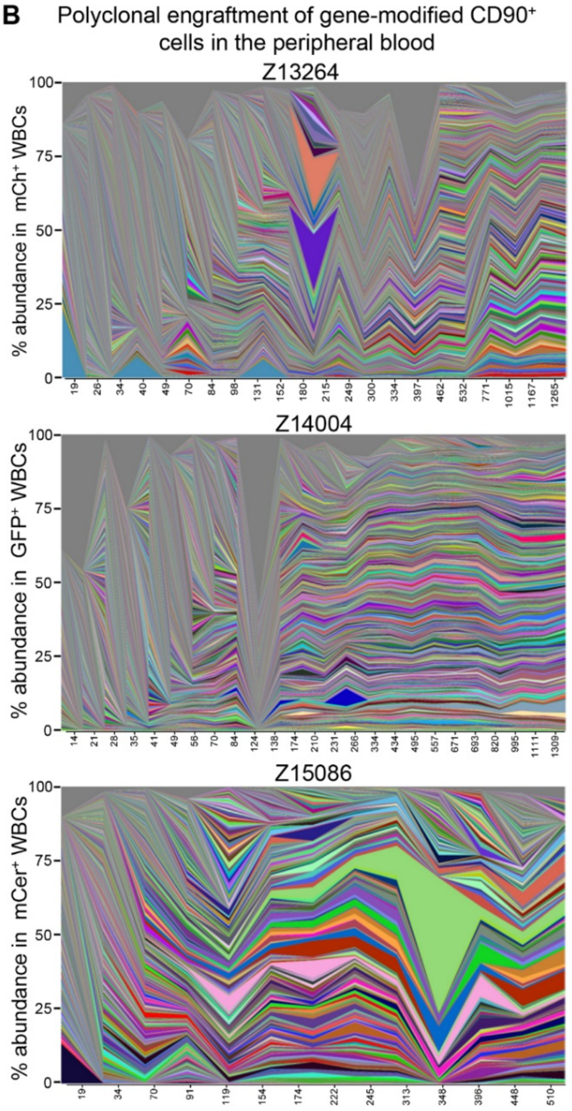


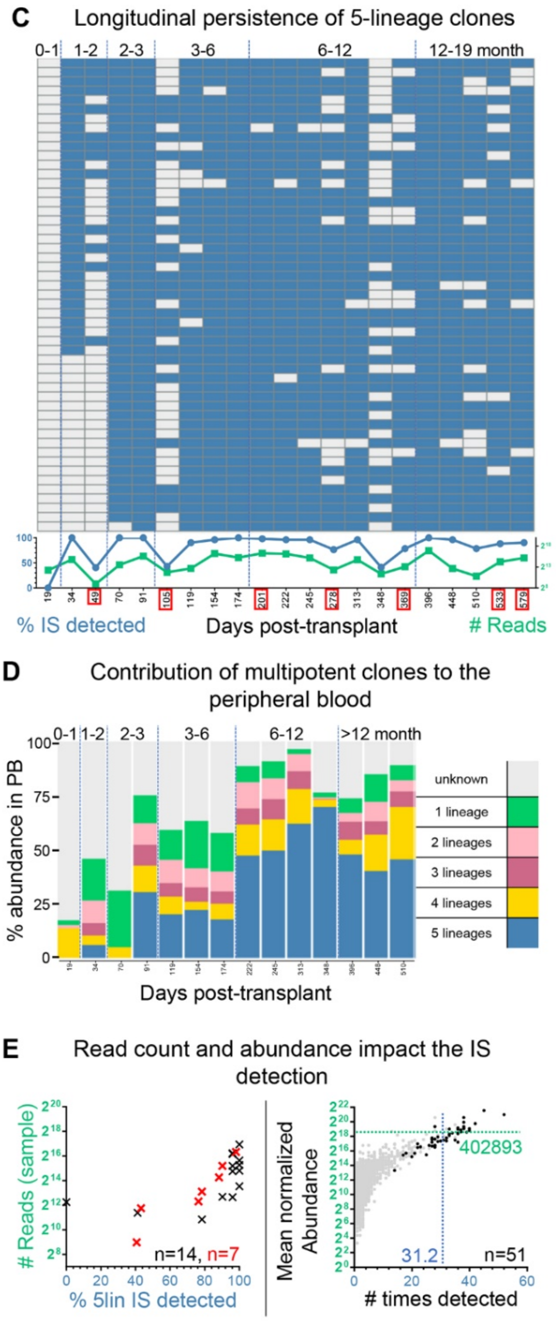


**Figure S2. CD90 animal characteristics, polyclonality in the PB, and clone tracking in animal Z15086. (A)** Summary of CD90 animal and data characteristics. **(B)** Polyclonality within gene-marked WBCs in the peripheral blood and abundance of individual clones over time. **(C)** Longitudinal detection of multipotent 5-lineage clones. Blue indicates the presence, grey the absence of unique clones. Graph below heatmap: Frequency of 5-lineage clones detected (blue, left y-axis) and number of reads (green, right y-axis) over time. BM time points are highlighted in red. **(D)** Longitudinal contribution of 5-, 4-, 3-, 2-, and 1-lineage clones to the peripheral blood. Color code as defined in Figure 2A. **(E)** Impact of data quality and clonal abundance on the reliability of IS detection. Left graph: Correlation of the cumulative read count from a single PB (black symbol) or BM (red symbol) time point with the frequency of multipotent 5-lineage clones detected at the same time point. Right graph: Correlation of the mean normalized abundance of 5-lineage clones (black dots) with the number of times the clone was detected across all available samples. Grey background indicates clones associated with other groups. The mean normalized abundance and average number of times detected across all clones are indicated with the green horizontal line and the blue vertical line, respectively.

### Figure S3

**A** Integration site data summary for CD34 animals

| | J02370 | M05189 | T04228 | Z08103 | Z09132 |
|---|---|---|---|---|---|
| % Gene Marking | 21-56% | 5-15% | 17-33% | 1-2% | 11-24% |
| Follow-up [days] | 2574 | 767 | 1194 | 747 | 720 |
| # Samples | 21 | 3 | 6 | 18 | 16 |
| # Total IS | 15,808 | 3,852 | 10,227 | 6,624 | 9,969 |
| # IS in Gene | 7,765 | 1,747 | 4,469 | 3,176 | 4,368 |
| # Genes with IS | 4,521 | 1,494 | 3,187 | 2,368 | 3,260 |
| # of Sequencing Reads | 1,735,275 | 640,671 | 1,009,796 | 1,193,774 | 728,484 |

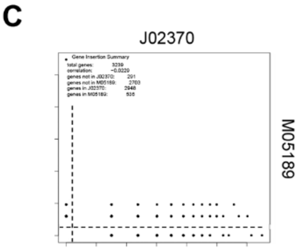


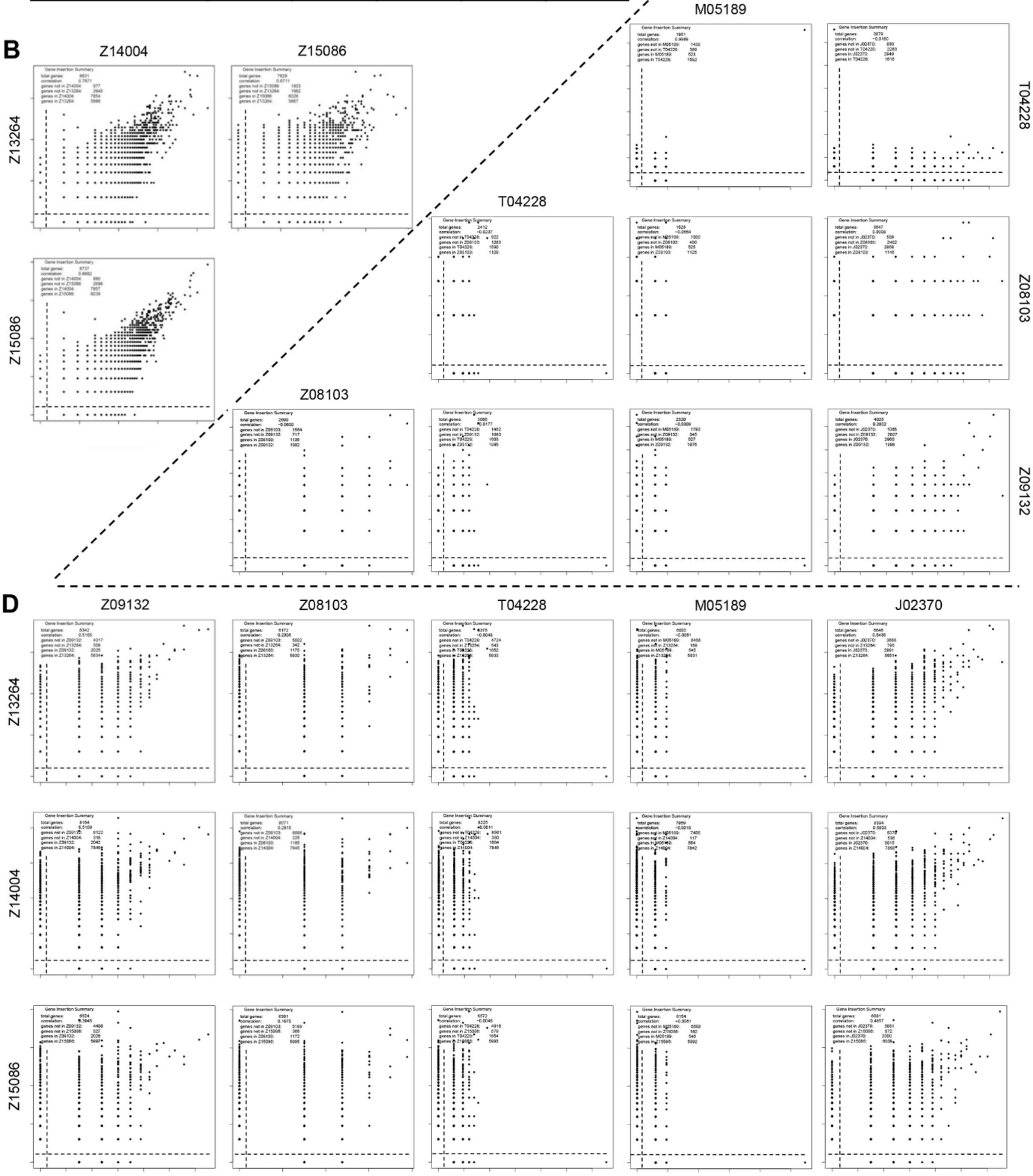


**Figure S3. Correlation of in-gene integration sites between CD34 and CD90 animals. (A)** Summary of CD34 animal and data characteristics. **(B-D)** Pair-wise correlation of the normalized abundance of in-gene IS across (B) CD90 animals, (C) across CD34 animals, and (D) of CD34 vs. CD90 animals. For genes with multiple unique IS, normalized counts were combined.

Figure S4

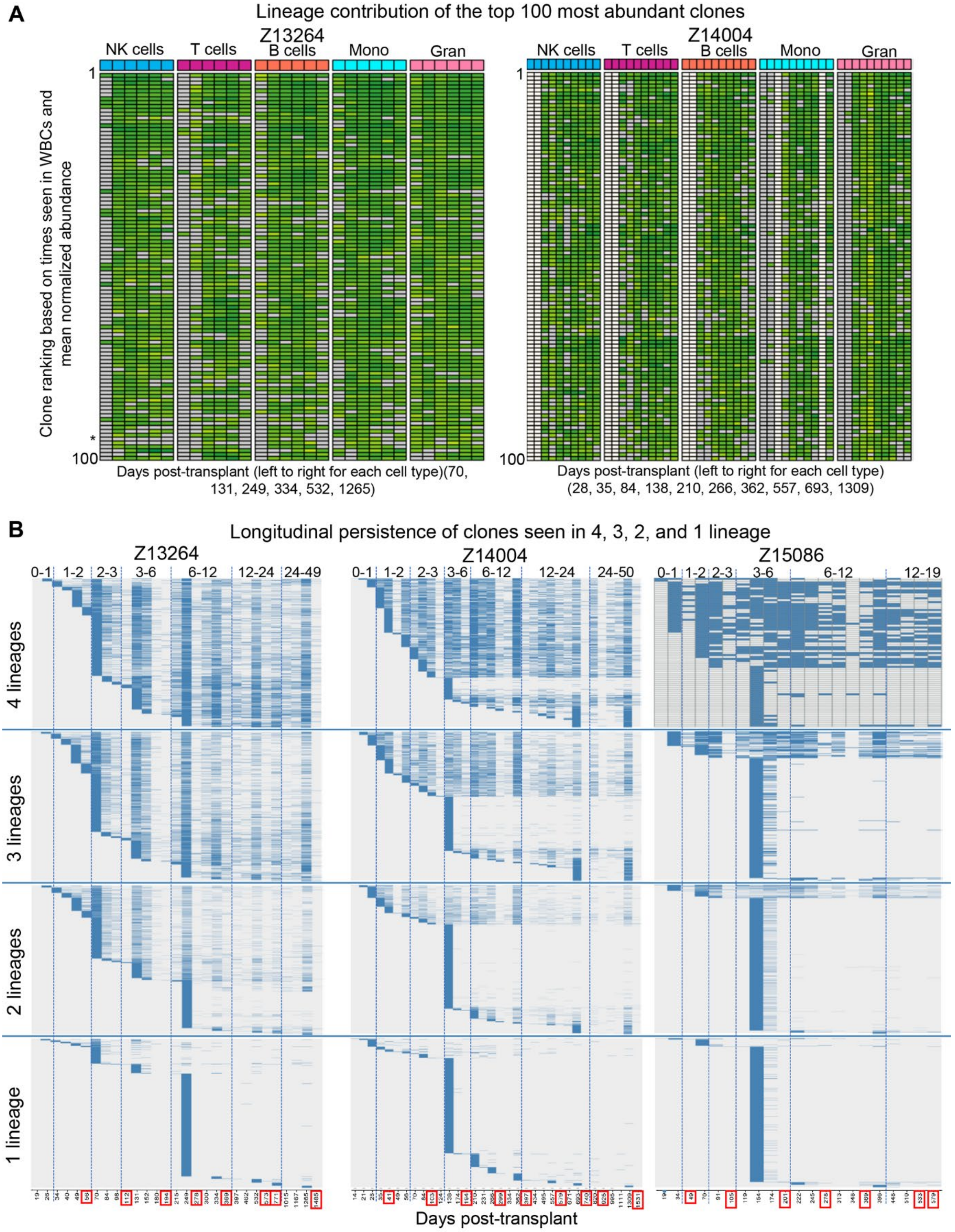


**Figure S4. Contribution and persistence of 4-, 3-, 2-, and 1-lineage clones. (A)** Impact of clonal abundance on the reliability of IS detection. Correlation of the mean normalized abundance of 4-, 3-, 2-, and 1-lineage clones (black dots) with the number of times the clone was detected across all available samples. Grey background indicates clones associated with other groups. The mean normalized abundance and average number of times detected across all clones are indicated with the green horizontal line and the blue vertical line, respectively. **(B)** Longitudinal detection of 4-, 3-, 2-, and 1-lineage clones. Blue indicates the presence, grey the absence of unique clones. BM time points are highlighted in red.

### Figure S5

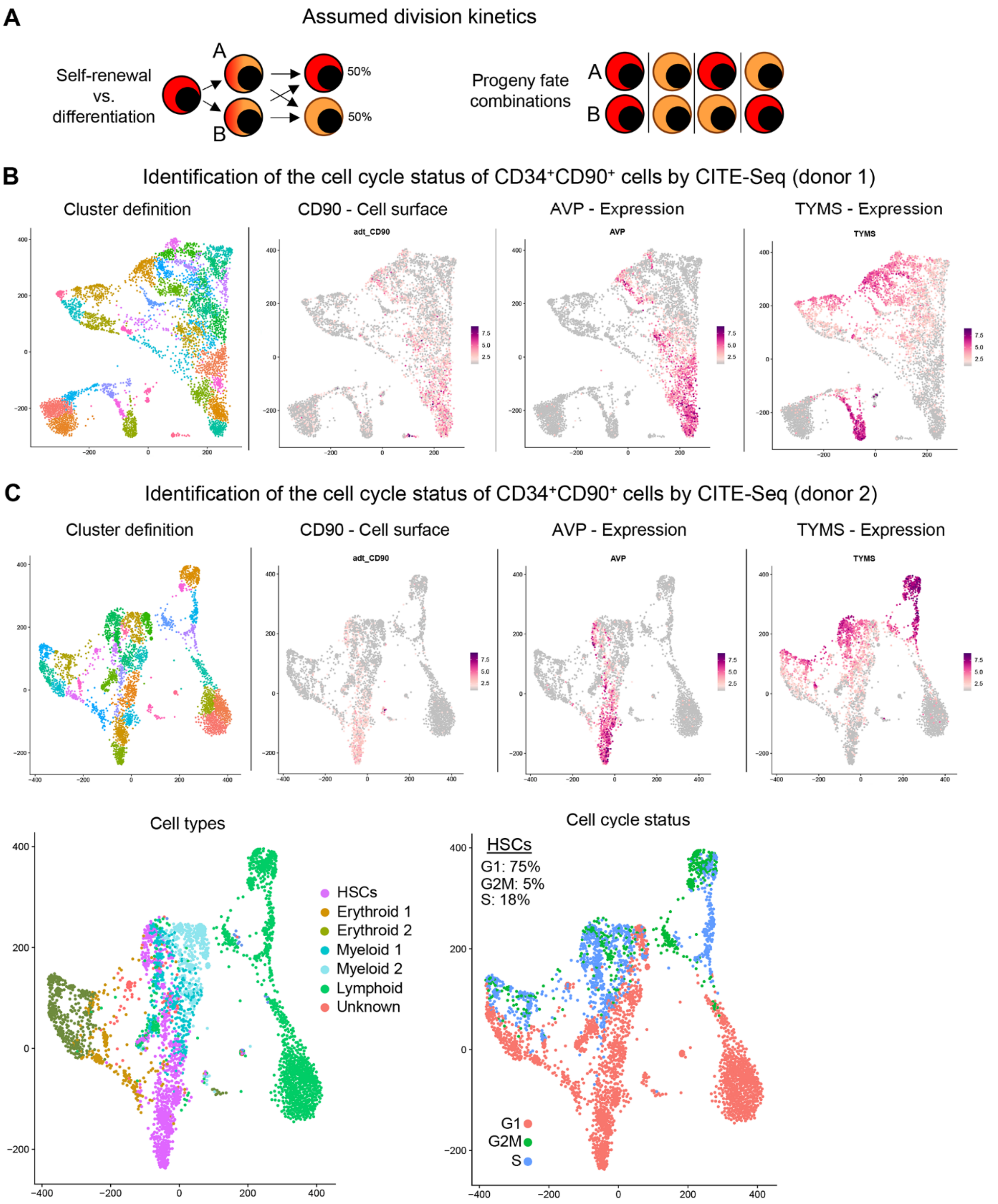


**Figure S5. Determination of parameters for the simulation of a stochastic HSC engraftment. (A)** Schematic of the assumed population neutral division of HSCs. Asymmetric and symmetric division of HSCs are expected to yield a 50%:50% chance for daughter cells after division to either maintain HSC potential or differentiate. **(B)** CITE-Seq of a representative human steady-state bone marrow CD34$^+$ population. RNAseq data was analyzed using Seurat to identify clusters (right top graph) of transcriptionally-distinct subsets. Identification of the HSC-enriched CD34$^+$CD90$^+$ subset was achieved overlaying the cell surface expression of CD90 (bottom left graph) and the expression of AVP (right top graph). Finally, TYMS expression, a representative cell cycle-associated gene, was overlaid to determine proliferating and quiescent CD34$^+$CD90$^+$ cells (bottom right graph).

Figure S6

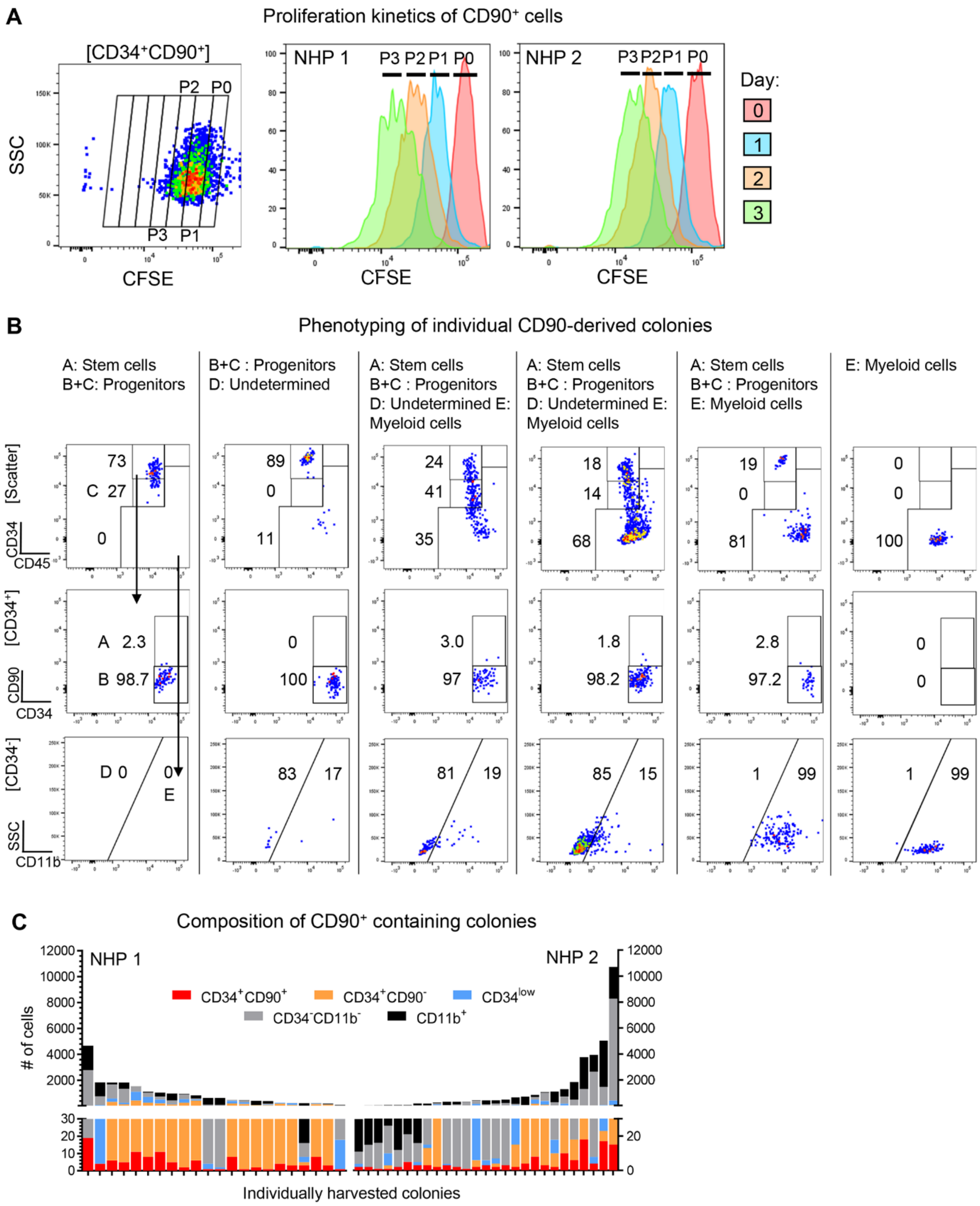


**Figure S6. Proliferation and differentiation of CD34$^+$CD90$^+$ cells in CFC assays. (A)** Representative flow cytometric assessment of CFSE staining of CD34$^+$CD90$^+$ cells on day 1. Histogram overlay of CFSE staining on the day before culture (day 0, red), at day 1 (blue), day 2 (orange), and day 3 (green). **(B)** Flow cytometric assessment of single CD34$^+$CD90$^+$-derived colonies. Cells were classified as A) CD34$^+$CD90$^+$CD45$^+$ HSCs, B+C) CD34$^+$CD90$^-$CD45$^+$ progenitors, D) CD34$^-$CD90$^-$CD11b$^-$CD45$^+$ undetermined/non-committed, and E) CD34$^-$CD90$^-$CD11b$^+$CD45$^+$ myeloid cells. **(C)** Size and composition of CD34$^+$CD90$^+$-containing colonies after 11 days of culture. Phenotypes and cell numbers were determined by flow cytometry. Phenotypically-defined stages are colored as indicated in the legend in the middle of the graph.
